# Zebrafish Functional Screening of FDA-Approved Drugs for Autosomal Dominant Retinitis Pigmentosa Caused by RHODOPSIN Q344X Mutation

**DOI:** 10.64898/2026.04.18.719270

**Authors:** Beichen Wang, Logan Ganzen, Emre Coskun, Rebecca James, Truc Kha, Xiaoguang Zhu, Jillian Alice New, Motokazu Tsujikawa, Yuk Fai Leung

## Abstract

Retinitis Pigmentosa (RP) is a group of inherited retinal degenerations for which most subtypes lack effective drug treatments. This challenge is particularly critical for autosomal dominant (ad) RP, which is often unsuitable for gene replacement therapy. To address this challenge, we screened an FDA-approved compound library using a zebrafish adRP model expressing a human RHODOPSIN transgene with the Q344X mutation. The screen evaluated drug effects on larval visual behavior by assessing the visual-motor response (VMR). Four compounds significantly improved VMR in Q344X zebrafish: amitriptyline, difluprednate, maprotiline, and prednisolone. Further characterization revealed that these hits act through distinct mechanisms, including reducing rod death, promoting rod neogenesis, and enhancing the function of extraocular photoreceptors. Together, these findings demonstrate the potential to repurpose these drugs for adRP caused by the RHO Q344X mutation, providing preclinical candidates and revealing potential targets for future drug development.

## INTRODUCTION

Retinitis Pigmentosa (RP) comprises a group of genetic disorders that cause retinal degeneration, affecting approximately 1.5 million individuals worldwide.^1–3^ RP results from mutations in more than 260 genes.^4^ These mutations are inherited in autosomal dominant (AD), autosomal recessive (AR), or X-linked patterns, primarily leading to the degeneration of rod photoreceptors and, in some cases, cone photoreceptors as well.^4–6^ In affected patients, rod degeneration manifests as clinical symptoms including night blindness and loss of peripheral vision, which may progress to complete blindness in severe cases.^6,7^ Moreover, these patients frequently endure socioeconomic hardships and psychological stresses, significantly diminishing their quality of life.^8–10^ However, effective treatments for RP remain scarce. While many are undergoing preclinical development and clinical trials, few ultimately advance to clinical use.

Current therapeutic approaches for RP, both approved and investigational, focus on correcting the underlying genetic mutations or addressing their downstream consequences. The former approach receives significant attention because it may cure RP, and implementation depends on the mode of inheritance.^11–13^ For instance, arRP and many X-linked RPs are characterized by a loss of normal gene function; they can be treated by gene supplementation therapy. A successful example is Luxturna, an FDA-approved gene therapy that delivers a functional *RPE65* gene to patients suffering from Leber congenital amaurosis caused by biallelic loss-of-function mutations in the *RPE65* gene.^14,15^ In adRP, however, gene mutations lead to a gain of abnormal protein function. Hence, these genes must be silenced by either RNAi or CRISPR/Cas gene editing before gene supplementation.^11,16^ Newer CRISPR/Cas variants that enable base editing also hold great promise for correction of mutations across all modes of inheritance.^17–19^ Although gene therapy may offer a permanent cure for RP, it is largely gene-specific and is more suitable for early-stage patients before significant photoreceptor degeneration has occurred.^11^ In addition to gene therapy, other treatments address the downstream consequences of genetic mutations, including stem cell therapy, vision restoration technologies, and pharmacological modulation. Stem-cell therapy involves transplanting differentiated retinal cells or their precursors into the damaged retina to preserve residual function or restore lost vision.^4,20^ Vision restoration, by contrast, aims to augment degenerating photoreceptors or bypass them altogether through visual prosthetics and optogenetics.^21^ Finally, pharmacological modulation utilizes pharmacological agents to influence relevant biological processes, including preventing cell death^22^, or restoring visual function^23,24^. Although pharmacological agents do not permanently correct underlying genetic defects and require repeated administration, they may slow disease progression and preserve vision.^25^ They are less invasive and often preferred by patients.^26^ These agents may target common biological processes conserved across different RP subtypes, irrespective of the specific mutation, thereby providing mutation-independent therapeutic effects.^25,27^ Consequently, pharmacological agents offer a unique and versatile treatment option for RP and allow for synergy with other treatment approaches.^28,29^

New pharmacological agents for RP are typically identified through either hypothesis-driven or exploratory strategies. In the hypothesis-driven strategy, potential drug targets are identified through prior dissection of a biological mechanism or pathway. These targets guide the design of precise perturbation approaches. For example, activation of metabolism in photoreceptors has been observed as a viable therapeutic approach.^30–33^ This insight has led to the identification and characterization of drugs targeting specific proteins involved in the process. In the exploratory strategy, drug targets are identified through screening compounds on disease models. The compounds can either be selected for their effects on potential disease mechanisms or from a library with diverse chemical structures and activities. The latter selection does not rely on prior knowledge of disease mechanisms or pathways, which are often unknown. It instead relies on carefully designed readouts in high-throughput screening (HTS) to identify drug hits that produce desirable outcomes in RP models, either *in vitro* or *in vivo*.^34,35^

*In vitro* HTS identifies hits based on improvements in biochemical assays or cellular behaviors in culture. For example, two recent *in vitro* HTS studies identified new drug hits using cell lines expressing the mouse Rhodopsin protein carrying a P23H mutation.^35,37^ One study screened 79,080 compounds and identified a new chemical chaperone that promoted the escape of the misfolded Rhodopsin protein from the endoplasmic reticulum and enhanced its stability.^35^ Using the same model, the other screened 68,979 compounds and identified five hits that promoted degradation of the misfolded Rhodopsin protein to mitigate its toxic effects on rods.^37^ Even though *in vitro* HTS is efficient, consistent, and amenable to larger-scale screening than its *in vivo* counterparts, it relies on biochemical assays and cell culture systems whose findings will have to be validated by *in vivo* models.^38^ Alternatively, *in vivo* HTS can directly identify these benefits by detecting drug effects on phenotypic, physiological, or functional improvements in RP animal models, thereby potentially reducing the attrition in the drug development pipeline.^39,40^ For instance, a phenotypic HTS evaluated the neuroprotective effects of 2,934 primarily FDA-approved compounds in a zebrafish model with inducible rod ablation, as measured by an increase in rod-reporter fluorescence.^41^ Among the 11 leads identified, three also promoted photoreceptor survival in retinal explants from *rd1* mice. Our group has long been developing a functional platform to evaluate drug effects in zebrafish models of retinal degeneration.^42–45^ This platform measures the visual-motor response (VMR) of zebrafish larvae, a startle response exhibited during sudden light changes.^42^ It was first used to assess visual function under photopic stimulation^46^. We then modified it to measure scotopic VMR mediated by rod functions^45^. This modification enabled us to quantify behavioral deficits due to rod degeneration and to screen drugs for RP using functional readouts.^45,47^ For instance, we screened an antioxidative compound library of 84 compounds on a zebrafish RP model carrying a human *RHODOPSIN* transgene with a Q344X mutation.^45,48^ We identified carvedilol, an FDA-approved drug for cardiovascular disease, which improved the VMR and increased rod number in the Q344X larvae ^45^

In this study, we extended our platform and screened an FDA-approved compound library of 1,430 compounds, reasoning that the identified hits could be repurposed for treating RP, an approach that potentially delivers new drugs faster and at lower cost than the traditional *de novo* approach.^49,50^ Our screen identified four novel hits that improved the VMR of the Q344X larvae. We also conducted initial functional analyses of these four hits to facilitate their future characterizations.

## RESULTS

### Screening of FDA-approved compounds in Q344X RP zebrafish via a visual motor response (VMR) assay

To identify compounds that improve visual function in Q344X RP larvae, a VMR screen was conducted using an FDA-approved library of 1,430 compounds (Figure 1A). Prior to screening, 191 compounds (13.4%) were excluded due to lethality or developmental toxicity observed in zebrafish larvae at 10 µM. The remaining 1,239 non-toxic compounds were screened in the VMR assay, with each compound tested on 24 Q344X larvae (Figure 1B). Treatment at 10 µM began at 5 days post-fertilization (dpf), and VMRs were measured at 7 dpf. The control groups were WT and Q344X larvae treated with equal volumes of drug carriers (DMSO or water) following the same treatment scheme, and their VMR data were obtained from our previous screen.^45^ After the drug screening, the positive hits were identified by criteria based on two prior observations in VMR. First, the control Q344X larvae showed a significant reduction in average total distance traveled after light offset compared to wildtype (WT) larvae, particularly during the first second (Supp Figure 1), attributed to rod degeneration in Q344X larvae.^45,48^ This observation revealed that the VMR difference is the most pronounced at the first second between WT and Q344X larvae, and the difference at the initial second was primarily driven by the retinal function.^45,47,51,52^ Second, the compound carvedilol was previously identified as a hit that improved Q344X VMR during the first 30 seconds after light offset, but not the first second.^45^ Given these two observations, two criteria were defined to identify positive hits in this screening of FDA-approved compounds: (1) compounds that increased the average total distance traveled by Q344X larvae to a similar level as WT larvae during the first 30 seconds following light offset, designated as Type I hits (Figure 1C); and (2) compounds that significantly increased the total distance travelled by Q344X larvae traveled compared with Q344X controls in the first second following light offset, designated as Type II hits (Figure 1D). Type I and Type II hits were subsequently identified through clustering analysis and hypothesis testing, respectively.

**Figure 1.**
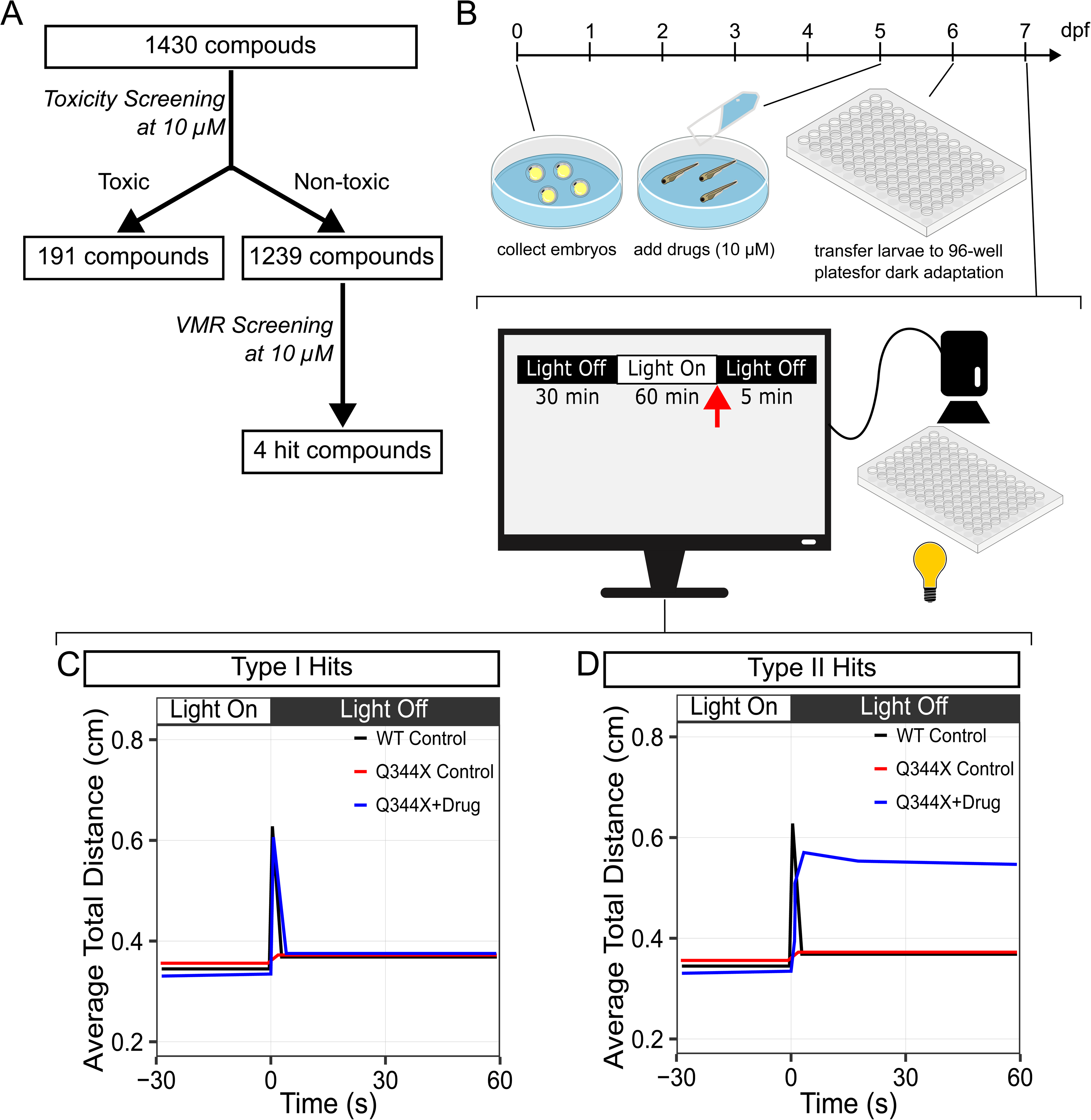
Design and setup for screening FDA-approved compounds via zebrafish visual-motor response (VMR). **(A**) Compound selection from the SelleckChem FDA-approved library for VMR screening. The FDA-approved library contains 1,430 compounds. A total of 191 compounds were excluded due to their observed toxicity in zebrafish larvae between 5 and 7 dpf at 10 µM. The remaining 1,239 compounds were subsequently screened in the VMR assays using the same treatment duration and concentration. **(B)** Workflow for compound treatment and VMR screening in zebrafish larvae. Embryos were collected and maintained in E3 medium until 5 dpf. At 5 dpf, compounds were added to the E3 medium at a final concentration of 10 µM. At 6 dpf, larvae were transferred to 96-well plates and placed in light-shielded boxes for dark habituation. At 7 dpf, the habituated larvae were placed in the VMR machine, which recorded their behavioral responses to light stimuli. The VMR protocol contains a 30-minute dark period (light-off), a 1-hour light period (light-on), and a 5-minute light-off period. **(C and D)** Illustrations of VMR screening results and criteria for hit identification. Light-off VMR is quantified by the average total distance traveled by larvae. Here, Type I and Type II hits are defined as follows: Type I hits restore Q344X VMR to WT control levels during the first 30 seconds after light offset; Type II hits elevate Q344X VMR above Q344X control levels during the first second after light offset. In these illustrations, white and black boxes above the plots indicate the light-on and light-off conditions during the time course, respectively.

Clustering analysis was first applied to the VMR data to identify Type I hits that improved the average total distance traveled by Q344X larvae to a level comparable to that by WT controls during the first 30 seconds following light offset. To minimize bias during hit identification, three clustering algorithms were employed: hierarchical clustering (HC),^53^ k-means clustering (k-means),^54^ and Gaussian Mixture Model with Expectation Maximization (GMM-EM).^55^ Clustering results from all three methods were visualized using Uniform Manifold Approximation and Projection (UMAP) (Figure 2A and Supp Figure 2), which projected high-dimensional VMR data into two-dimensional space.^56^ Aggregation of the results of three clustering algorithms yields 25 unique Type I hits from the 1,239 non-toxic compounds (Figure 2B). These 25 hits increased the average total distance traveled by Q344X larvae to the levels observed in WT larvae during the first 30 seconds following light offset.

**Figure 2.**
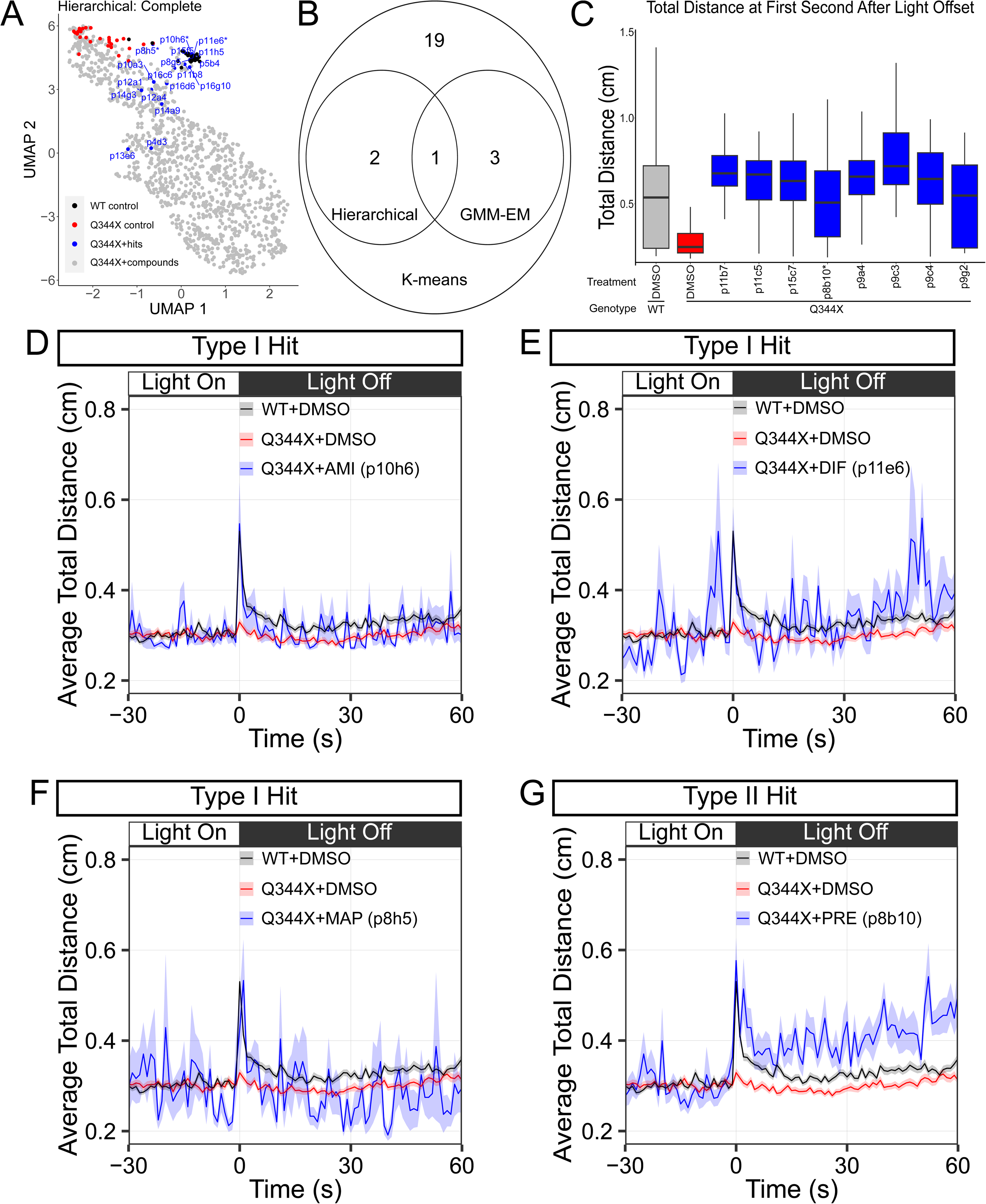
Hit identification by clustering analysis and hypothesis testing. **(A)** Identification of Type I hits using hierarchical clustering with complete linkage, visualized by a UMAP plot. Clusters were assigned by cutting the dendrogram at the level that partitioned the WT and Q344X groups. On the UMAP plot, each dot represents the projection of average total distance curves during the first 30 seconds following light offset for each treatment group. The color of the dot represents a specific genotype and its treatment outcome: WT control (black), Q344X control (red), drug-treated Q344X groups clustered with WT control (Q344X+hits; blue), and drug-treated Q344X groups that did not cluster with either WT or Q344X control groups (Q344X+compounds; grey). The codenames of the Type I hits are labeled next to the Q344X groups treated with these compounds. Additional results generated by other clustering algorithms are presented in Supp Figure 2. **(B)** A Venn diagram showing the number of Type I hits identified by all three clustering algorithms: hierarchical clustering (Hierarchical), k-means clustering (K-means), and Gaussian Mixture Model with Expectation Maximization (GMM-EM). **(C)** A box-and-whisker plot of the total distance travelled by control groups and Q344X groups treated with Type II hits during the first second following light offset. **(D–G)** Average total distance curves of the 4 hit-treated Q344X groups (blue) and the control groups (WT+DMSO, black; Q344X+DMSO, red) during light offset. These 4 compounds demonstrated consistent positive effects in both the initial screening and confirmatory testing. The confirmatory testing results are presented in Figure 3. Sample size: WT+DMSO and Q344X+DMSO, n = 48 larvae per genotype × 9 biological replicates; Q344X+AMI, n = 24 larvae; Q344X+DIF, n = 24 larvae; Q344X+MAP, n = 24 larvae; Q344X+PRE, n = 24 larvae. In these plots, the ribbons indicate the standard error of the mean. The white and black bars above the plots denote the light-on and light-off phases of the time course, respectively.

Second, hypothesis testing was used to identify Type II hits that significantly increased the total distance traveled by Q344X larvae compared with that by Q344X controls during the first second following light offset. The Welch two-sample test was employed, and multiple hypothesis testing was corrected using the Bonferroni method. Eight Type II hits were identified from 1,239 non-toxic drugs (adjusted p-value < 0.05; Figure 2C).

A total of 34 hits of both types were identified in the initial screening. These hits were subsequently validated through confirmatory VMR testing using compounds obtained from alternative vendors (Supp Table 1). Out of the 34 hits identified in the initial screening, 4 compounds in the confirmatory VMR testing significantly increased the average total distance traveled by Q344X larvae during the first second following light offset, compared with the control treatment. These 4 compounds were amitriptyline hydrochloride (AMI, codename p10h6; Type I hit), difluprednate (DIF, codename p11e6; Type I hit), maprotiline hydrochloride (MAP, codename p8h5; Type I hit), and prednisolone acetate (PRE, codename p8b10; Type II hit). Their initial screening profiles are shown in Figure 2D–G, their confirmatory VMR profiles are presented in Figure 3A–D, and the statistical testing results for these confirmatory VMR profiles are shown in Supp Table 2. These initial and confirmatory VMR results indicate that the 4 identified compounds have the greatest therapeutic potential to enhance visual function in Q344X RP larvae.

**Figure 3.**
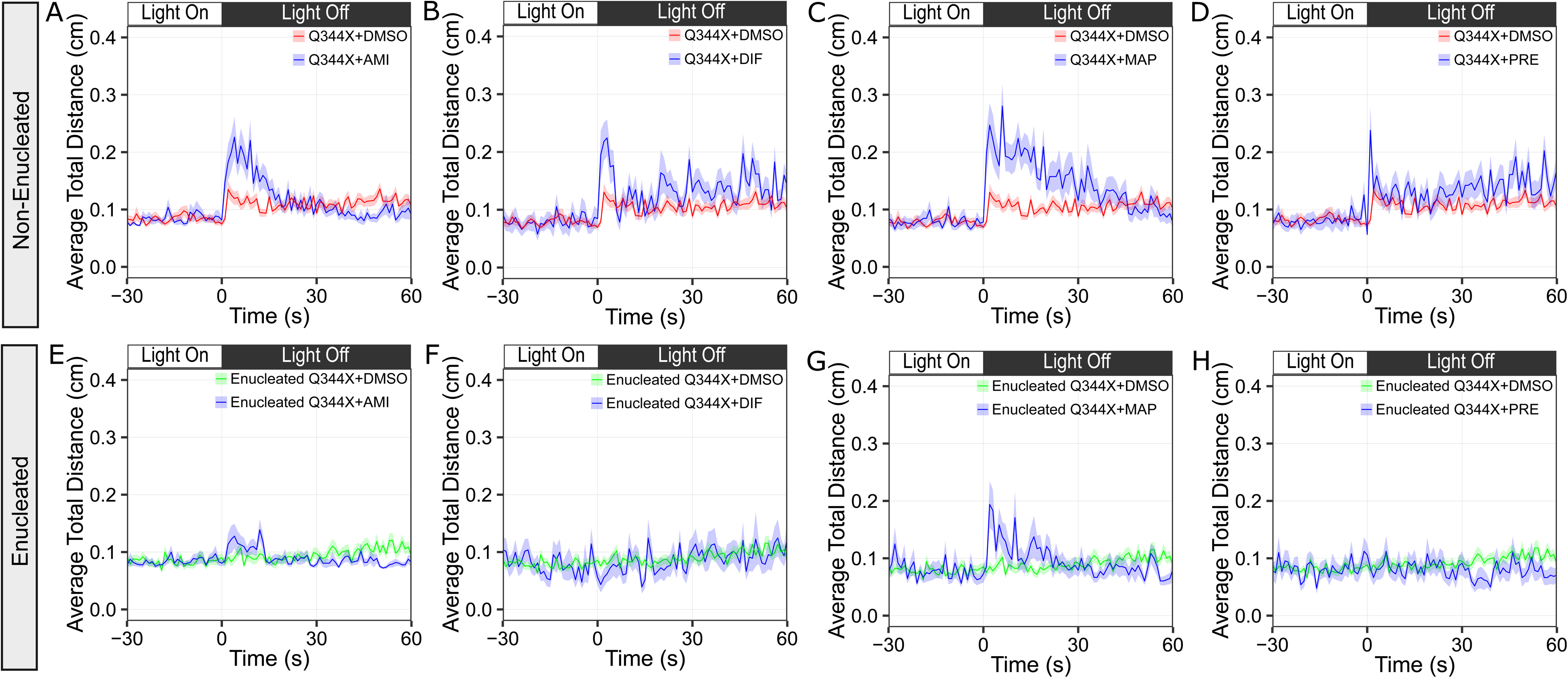
Average total distance curves of enucleated and non-enucleated Q344X larvae measured during the light offset period following treatment with hit compounds. **(A–D)** Average total distance curves of non-enucleated Q344X larvae treated with hit compounds (blue) or DMSO (red). **(E–F)** Average total distance curves of enucleated Q344X larvae treated with hit compounds (blue) or DMSO (green). In all plots, the ribbons indicate the standard error of the mean. The white and black bars above the plots denote the light-on and light-off phases of the time course, respectively. The corresponding sample size and statistical test results are provided in Supp Table 2.

### Investigating Hits Efficacy on Rod Function and Rod Number in Q344X RP Zebrafish

These 4 hits were subsequently evaluated for their effects on rod function and counts. Treatment with these hits might have enhanced VMR in Q344X larvae by improving rod function, increasing rod number, or enhancing the function of extraocular photoreceptors. VMR depends on retinal function^45–47^ as evidenced by the substantial reduction of VMR in enucleated Q344X and WT larvae (Supp Figure 3).^45^ However, extraocular function could also contribute to VMR.^57^ To assess the extent to which retinal and extraocular inputs contributed to the effects of hits, treatments were applied to enucleated Q344X larvae, and their light-off VMR was recorded. During the first second following light offset, enucleated Q344X larvae treated with AMI, DIF, or PRE showed no significant difference in average total distance traveled compared with enucleated larvae treated with DMSO (Figure 3E, F, and H, and Supp Table 2). Enucleated Q344X larvae treated with MAP did not show a significant increase in average total distance traveled during the first second following light offset (Figure 3G and Supp Table 2). However, these larvae showed a significant increase in average total distance traveled during the first 30 seconds after light offset, compared with enucleated larvae treated with DMSO (Hotelling’s T^2^ test, T^2^ = 1.4214, df_1_ = 31, df_2_ = 62.8159, p-value = 0.04373). These results suggested that the effects of AMI, DIF and PRE were not mediated through extraocular photoreceptors but were primarily mediated through the retina, thereby improving the Q344X VMR during the first second following light offset. In contrast, the effect of MAP was partially mediated through extraocular photoreceptors, contributing to the Q344X VMR during the later seconds after light offset.

These hits might have improved rod-driven VMR in Q344X larvae by enhancing residual rod function or increasing the number of rods. For instance, from 5 to 7 dpf, WT larvae exhibited significantly higher rod counts on each respective day (Supp Figure 4 and Supp Table 3), whereas Q344X larvae progressively lost their rods in the medial outer nuclear layer (ONL) (Supp Figure 4). To determine the extent to which the hits increased rod number, the *Tg(-3.7rho:EGFP)* reporter line,^58^ which expresses the enhanced green fluorescent protein (EGFP) in rods, was bred into the Q344X background, enabling fluorescent rod detection. Following the drug treatment initiated at 5 dpf, the rods were quantified in the Q344X retina at 7 dpf. At 7 dpf, Q344X larvae treated with AMI, MAP, or PRE exhibited higher rod counts compared with DMSO-treated Q344X larvae, particularly in the dorsolateral and ventrolateral ONL (Supp Figure 5B, D, and E), but the difference was not statistically significant (Figure 4A and Supp Table 4). Q344X larvae treated with DIF exhibited significantly higher rod counts compared with DMSO-treated Q344X larvae (Figure 4A and Supp Table 4). The extra rods were located in the dorsolateral, ventrolateral, and medial ONL (Supp Figure 5C). These results indicate that the improved VMR following DIF treatment might have resulted from an increase in rod number, whereas the improved VMR following treatment with AMI, MAP, and PRE was unlikely to arise from a change in rod number.

**Figure 4.**
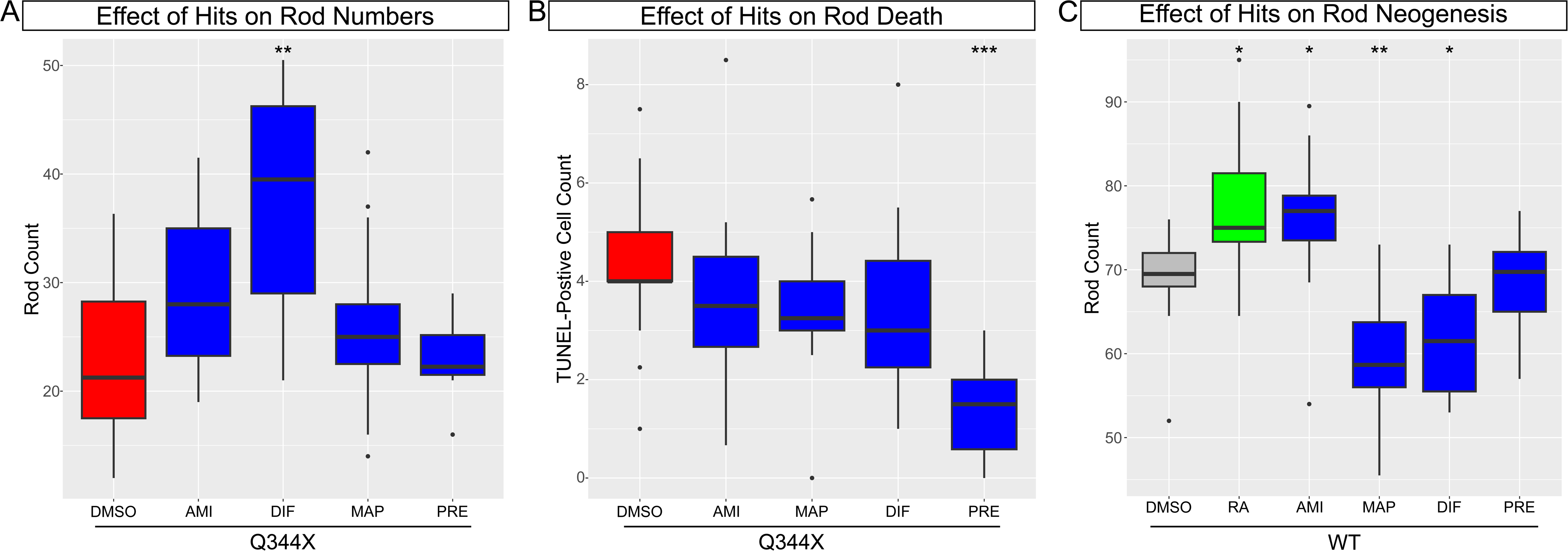
Histology quantification following treatment with hit compounds. **(A)** Box-and-whisker plots showing rod counts at 7 dpf in Q344X retinas following treatment with DMSO (red) or hit compounds (blue). Rods were visualized using the *Tg(-3.7rho:EGFP)* transgene. The corresponding sample size and statistical test results are provided in Supp Table 4. Representative images of retinal cryosections are provided in Supp Figure 5. **(B)** Box-and-whisker plots showing TUNEL-positive cell counts at 7 dpf in the outer nuclear layer of Q344X retinas following treatment with DMSO (red) or hit compounds (blue). TUNEL-positive cells were visualized on cryosections according to the manufacturer’s protocols. The corresponding sample size and statistical test results are provided in Supp Table 6. Representative images of retinal cryosections are provided in Supp Figure 7. **(C)** Box-and-whisker plots showing rod counts at 7 dpf in WT retinas following treatment with DMSO (grey), retinoic acid (RA, green), or hit compounds (blue). Hit compounds were applied at 10 µM, and RA at 1.2 µM. All treatments were administered from 5 to 7 dpf, consistent with the VMR screening protocol. The corresponding sample size and statistical test results are provided in Supp Table 7. Representative images of cryosections are provided in Supp Figure 8. Significance levels: *, p-value < 0.05; **, p-value < 0.01; ***, p-value < 0.001.

**Figure 5.**
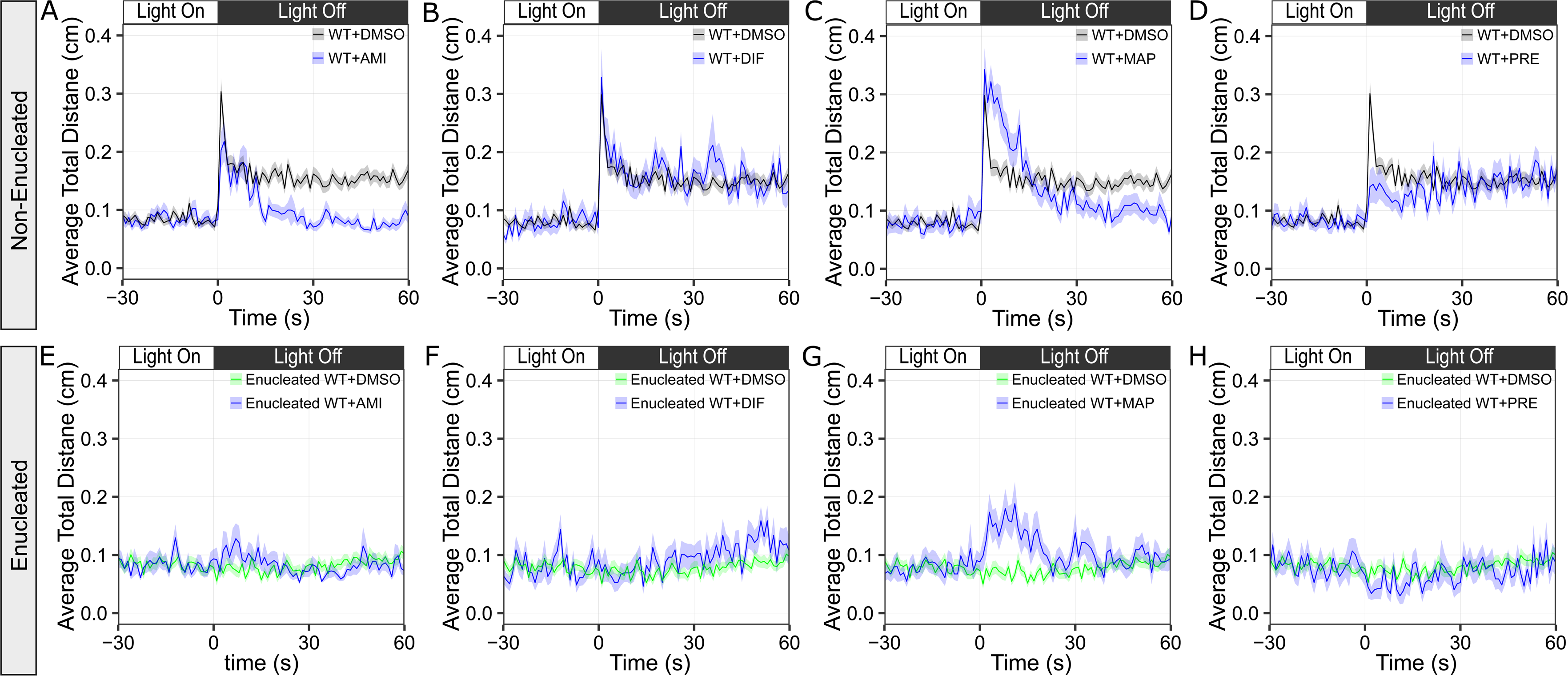
Average total distance curves of enucleated or non-enucleated WT larvae following treatment with hit compounds during light-offset. **(A–D)** Non-enucleated WT larvae treated with hit compounds (blue) or DMSO (black). **(E–F)** Enucleated WT larvae treated with hit compounds (blue) or DMSO. In all plots, the ribbons indicate the standard error of the mean. The white and black bars above the plots indicate the light-on and light-off phases of the time course, respectively. The corresponding sample size and statistical test results are provided in Supp Table 8.

### Mechanism of Rod Number Increase: Reduction in Cell Death

The increase in rod number following hit treatment might have resulted from reduced cell death or increased rod generation. First, cell death in drug-treated Q344X larvae was assessed using the terminal deoxynucleotidyl transferase dUTP nick-end labeling (TUNEL) assay, as the predominant mechanism of rod degeneration in RP involves apoptosis.^48,59,60^ The TUNEL-positive cells were quantified in the ONL of larval retinas. Between 5 and 7 dpf, WT larvae exhibited significantly fewer TUNEL-positive cells than Q344X larvae in the ONL on each corresponding day (Supp Figure 6 and Supp Table 5). Subsequently, TUNEL-positive cells in Q344X retinas were quantified at 7 dpf following hit compound treatment initiated at 5 dpf. At 7 dpf, Q344X retinas treated with AMI, DIF, or MAP exhibited lower TUNEL-positive cell counts compared with DMSO-treated controls, but the differences were not statistically significant (Figure 4B and Supp Table 6). In contrast, PRE-treated Q344X retinas exhibited a significantly lower TUNEL-positive cell count compared with DMSO-treated controls (Figure 4B and Supp Table 6). These results suggest that PRE treatment reduced rod death, while AMI, DIF, or MAP did not.

### Mechanism of Rod Number Increase: Rod Generation

In addition to reduced cell death, the increase in rod numbers might have resulted from enhanced rod generation following hit treatment. In larval retinas, new rods arise through regeneration or neogenesis.^61,62^ Although regeneration enhancement was a plausible mechanism, the duration required for drug-induced regeneration exceeded the treatment window used in this study.^63–65^ Therefore, the analysis focused on rod neogenesis as a possibility. To this end, a rod neogenesis assay was conducted in the WT larvae with the *Tg(-3.7rho:EGFP)* transgene. These larvae were treated with hit compounds and 1.2 µM retinoic acid (RA), which promotes rod generation,^66,67^ from 5 to 7 dpf. At 7 dpf, RA-treated retinas exhibited significantly higher rod counts compared with DMSO-treated control retinas (Figure 4C and Supp Table 7). These results demonstrated that the RA treatment in the current scheme was sufficient to promote rod neogenesis. AMI-treated WT retinas exhibited significantly higher rod counts compared with DMSO-treated retinas, and the values were comparable to those observed in RA-treated retinas (Figure 4C and Supp Table 7). The extra rods were located in the dorsolateral and ventrolateral ONL (Supp Figure 8C). WT retinas treated with DIF or MAP exhibited significantly reduced rod counts compared with DMSO-treated retinas (Figure 4C and Supp Table 7). PRE-treated retinas also exhibited reduced rod count compared with DMSO-treated retinas, but the reduction was not statistically significant (Figure 4C and Supp Table 7). These results suggest that AMI promoted rod neogenesis, MAP and DIF reduced rod abundance, and PRE had no effect on rod neogenesis.

### Validation of Altered Scotopic Retinal Function after Hit Treatment

Although AMI stimulated neogenesis, the total rod number was largely unchanged in Q344X retinas. In addition, AMI, MAP, and PRE significantly improved the scotopic VMR of Q344X larvae. These results suggest that visual recovery is not strictly mediated by an increase in the rod number in Q344X larvae, but may also result from an enhancement of the scotopic retinal function in Q344X larvae. To test this hypothesis, hit compounds were applied to the non-enucleated and enucleated WT larvae, and their VMR was evaluated by comparing the average total distance traveled by each group during the first second following light offset. First, non-enucleated WT larvae treated with AMI and PRE exhibited a significant reduction in the average total distance traveled compared with non-enucleated WT larvae treated with DMSO (Figure 5A & D, and Supp Table 8). In contrast, non-enucleated WT larvae treated with DIF and MAP exhibited an increase in average total distance traveled compared with non-enucleated WT larvae treated with DMSO, but the increase was not statistically significant (Figure 5B & C, and Supp Table 8). Second, enucleated WT larvae treated with AMI and DIF exhibited increased average total distance traveled during the first second following light offset compared with enucleated WT larvae treated with DMSO, but the increase was not statistically significant (Figure 5E–F and Supp Table 8). Enucleated WT larvae treated with MAP exhibited a significant increase in average total distance traveled during the first 30 seconds after light offset, compared with enucleated WT larvae treated with DMSO (Figure 5G; Hotelling’s T^2^ test, T^2^ = 2.3247, numerator df = 31, denominator df = 59.9136, p-value = 0.00013333). Enucleated WT larvae treated with PRE exhibited a reduction in the average total distance traveled during the first second following light offset compared with enucleated WT larvae treated with DMSO, but the reduction was not statistically significant (Figure 5H and Supp Table 8). Third, following treatment with any hits, enucleated WT larvae exhibited a significant reduction in the average total distance traveled compared with non-enucleated WT larvae (Supp Table 8). Altogether, these results suggest that DIF and MAP did not affect scotopic retinal function in WT larvae, whereas AMI and PRE reduced scotopic retinal function in WT larvae. These findings further suggest that MAP improved extraocular photoreceptor function, which partially contributed to improvement in scotopic VMR.

### Effects of Hit Compounds on Non-Visual System

Following the evaluation of the effects of hit compounds on rod function and number, their impact on non-visual sensorimotor responses was further assessed. To this end, the effect of each hit compound on a mechanosensory response was measured using a tapping assay.^68^ In this assay, larvae were placed in 96-well plates and tapped 11 times during a 2-minute recording using the VMR machine, operated under the same parameters as in previous experiments. In response to tapping, DMSO-treated WT and Q344X larvae exhibited no significant difference in average total distance traveled (Supp Figure 9A and Supp Table 9), suggesting the mechanosensory-motor circuitry remained unaltered in Q344X larvae. Q344X larvae treated with AMI or MAP exhibited a significant increase in average total distance traveled compared with DMSO-treated controls, whereas Q344X larvae treated with DIF or PRE exhibited no significant change. These results suggest AMI and MAP enhanced mechanosensory-motor function in the Q344X larvae, while DIF and PRE exerted no such effect.

### Summary of Identified Hits

Four hit compounds that improved Q344X scotopic VMR were identified, and their effects were characterized across multiple assays. A summary of these characterizations is provided in Table 1, together with the structural similarity among these compounds (Supp Figure 10). AMI and MAP exhibit structural similarity, as do DIF and PRE, suggesting that each pair might bind to common molecular targets and act through a common signaling pathway.

**Table 1.** Summary of hit compounds and their effects in Q344X and WT zebrafish. Abbr.: abbreviation; ↑: increase; ↓: decrease; NC: no change.

| Code Name | Drug Name | Abbr. | CAS | Hit Type | Q344X VMR | Enucleated Q344X VMR | Q344X Rod Number | Q344X TUNEL<sup>+</sup> Cell Number | WT Rod Number | WT VMR | Enucleated WT VMR | Mechanosensory Response |
| --- | --- | --- | --- | --- | --- | --- | --- | --- | --- | --- | --- | --- |
| p10h6 | Amitriptyline hydrochloride | AMI | 549-18-8 | I | ↑ | NC | NC | NC | ↑ | ↓ | NC | ↑ |
| p11e6 | Difluprednate | DIF | 23674-86-4 | I | ↑ | NC | ↑ | NC | ↓ | NC | NC | NC |
| p8h5 | Maprotiline hydrochloride | MAP | 10347-81-6 | I | ↑ | ↑* | NC | NC | ↓ | NC | ↑* | ↑ |
| p8b10 | Prednisolone acetate | PRE | 52-21-1 | II | ↑ | NC | NC | ↓ | NC | ↓ | NC | NC |

## DISCUSSION

Most RP subtypes currently lack effective treatments or approved therapies. To address this issue, this study aimed to repurpose FDA-approved drugs for adRP by screening a library of 1,430 FDA-approved compounds using a zebrafish adRP model carrying the human RHO Q344X mutation.^48^ This screening utilized VMR, a functional assay, to assess the drug efficacy in improving visual performance.^45,46^ The screening identified four FDA-approved drugs—AMI, DIF, MAP, and PRE—that improved the visual performance in Q344X zebrafish, providing novel hits for the treatment of Q344X adRP.

The four functional hits identified in this study added new pharmacological candidates for treating Q344X RP. Three Type I hits—AMI, DIF, and MAP—restored the Q344X VMR to the WT level (Figure 2D–F), while one Type II hit, PRE, improved the Q344X VMR (Figure 2G). Among the three Type I hits, AMI was consistently identified by all three clustering algorithms (Figure 2A–B and Supp Figure 2), indicating its robust capacity to rescue the Q344X VMR to the WT level. In addition to improving Q344X VMR and visual functions, the identified hits also exerted distinct effects on rod histology and functions. AMI promoted rod neogenesis (Figure 4C) but reduced scotopic retinal function in the WT retina (Figure 5A). DIF increased rod number in the Q344X retina (Figure 4A) but reduced it in the WT retina (Figure 4C). MAP reduced rod number in the WT retina (Figure 4C) and enhanced extraocular photoreceptor function (Figure 3G and Figure 5G). PRE reduced rod death in Q344X larvae (Figure 4B) while reducing the scotopic retinal function in the WT retina (Figure 5D). Although reduced cell death typically increases total cell number, PRE decreased rod death without changing the total rod number in Q344X larvae (Figure 4A). This finding suggests that PRE could have increased the ratio of functional rods within the Q344X ONL. In other words, the observed GFP-positive rods likely comprise a mixed pool of dying rods and functional cells, so reducing rod death could have shifted the population toward functionality without changing the total number. This shift corresponded with the increased scotopic VMR (Figure 2G and Figure 3D). Future studies should precisely label these rod populations to clarify the specific effect of PRE.

These results indicated that identified hits may exert their effect via distinct biological pathways. Based on the known indications and structural similarities (Supp Figure 10), these hits fall into two classes: antidepressants (AMI and MAP) and corticosteroids (DIF and PRE). These drug classes differ in their effects on non-visual function. Antidepressants enhanced mechanosensory responses (Supp Figure 9B and D), whereas corticosteroids did not (Supp Figure 9C and E). Nonetheless, the results suggested two plausible therapeutic targets in Q344X RP: 1) receptors targeted by AMI and MAP, including serotonergic, adrenergic, and histaminergic receptors;^69^ and 2) the glucocorticoid receptor (GR) targeted by DIF and PRE.^70,71^

Besides identifying novel hits, this study demonstrates the benefit of clustering algorithms in VMR analysis. Although hypothesis testing remains a standard analysis for screening data,^72^ clustering offers a complementary and unique perspective on drug efficacy in behavior screening. Clustering captures similarities and dynamics in behavior trajectories that may be overlooked by conventional hypothesis testing.^73,74^ For instance, clustering uniquely identified three Type I hits that hypothesis testing failed to identify, even during the first second after light offset, when the WT and the Q344X larvae differed the most (Figure 2 and Supp Figure 1). This result underscores the risk of relying solely on hypothesis testing for hit identification. Integrating hypothesis testing and clustering provides a more comprehensive approach to identifying hits and uncovering novel mechanisms.

Despite its contributions, this study has three limitations. First, the screening was conducted at a single concentration (10 µM) and treatment period (5 to 7 dpf) per compound. This design may overlook compounds that exhibit optimal efficacy at different concentrations or treatment periods, potentially explaining the divergent effects observed during characterization. Expanding the concentration and treatment period range will increase the number of tests, thereby posing a challenge to the current throughput of *in vivo* screening. In addition to increasing the number of screening machines, which can be costly, future VMR screening could implement an orthogonal pooling strategy to eliminate negatives more efficiently,^75^ thereby balancing the screening cost and efficiency. Second, our hit identification relied on two analytic criteria. Incorporating additional criteria could reveal more hits that partially rescue the Q344X VMR (gray points, Figure 2A, Supp Figure 2). For instance, our previous VMR screening identified carvedilol, which improved the Q344X VMR during the first 30 seconds after light offset but not in the first second.^45^ This finding suggests that effective compounds may improve the Q344X VMR with varying temporal dynamics. Future analysis could further examine these dynamics, utilizing feature-selection methods^51^ to identify time points that reveal meaningful behavioral differences induced by the drug, thereby facilitating more effective hit identification and prioritizing candidate compounds for downstream development. Third, the hit effect on visual function was assessed only at the retinal level but not at the rod level. Although scotopic VMR was specifically mediated by rod function^45,47^, the enucleation isolated the extraocular inputs from the retinal inputs but did not specifically investigate rod inputs. Future studies could measure electroretinogram responses^47^ or image glutamate release from rods in zebrafish larvae^76^ during scotopic light changes, thereby pinpointing the specific effect on rods.

In conclusion, this study identified and characterized four FDA-approved compounds that improved the visual function in the Q344X RP zebrafish, expanding the preclinical candidate pool for adRP. Since these four compounds are already approved by the FDA for human use, repurposing them may accelerate therapeutic development. Previous investigations also support their therapeutic potential: AMI was shown to protect rods from degeneration by upregulating glial cell line-derived neurotrophic factor;^77–80^ corticosteroids have been reported to improve photoreceptor health via GR agonism.^81–83^ Altogether, previous findings and our results build a compelling case for continued investigation and development of these four compounds. Further development of these pharmacological agents could offer affordable, accessible, and non-invasive treatment options for RP.

## METHODS

### Zebrafish Husbandry

The transgenic line *Tg(rho:Hsa.RH1_Q344X)* was generated and used in our previous studies,^45,48^ while *Tg(-3.7rho:EGFP)* was generated by others ^58^ and used in our studies.^45,48^ All animals were maintained at 28°C with a 14-h light/ 10-h dark cycle following standard procedures.^84,85^ To collect embryos, adults were placed sex-separated in breeding tanks the night before. They were mixed and bred from 9 a.m. to 11 a.m. the following day. Embryos were collected and then raised in the E3 medium in an incubator. The E3 medium was changed every day, and unhealthy embryos were discarded. All protocols were approved by the Purdue University Institutional Animal Care and Use Committee.

### Compounds and Treatment

Our study screened a Selleckchem FDA-approved compound library with 1430 compounds (Houston, TX). The validation experiments used compounds purchased from SelleckChem (Houston, TX), MedChemExpress (Monmouth Junction, NJ), and Sigma Millipore (St. Louis, MO). Detailed compound information is available in Supp Table 1. All compounds were prepared in 15 mL of E3 medium at a final concentration of 10 µM, a common starting concentration for screening in the zebrafish model.^86^ Each working solution was administered to 30 larvae per group from 5 to 7 dpf. Each group was kept in a single 100 × 15 mm Petri dish (DOT Scientific, Burton, MI) from 5 dpf. In the VMR experiment, larvae were transferred to 96-well plates with the working solution at 6 dpf and maintained there until 7 dpf (See next section). In all other experiments, larvae remained in the Petri dish with the working solution throughout the exposure period. This treatment period was chosen to maximize rod protection, as we previously observed a significant loss of Q344X rods starting at 5 dpf. At 7 dpf, most rods were degenerated, and the VMR displayed by the Q344X was significantly reduced compared with that displayed by the wildtype.^45,48^ The treatment was not refreshed during the experiment. For controls, an equal amount of compound carrier (DMSO or water) was added to the E3 medium. The final DMSO concentration was 0.1%.

### Visual-Motor Response Assay

At 6 dpf, drug-treated and control zebrafish larvae were transferred to Whatman UNIPlate square 96-well plates (VWR, Radnor, PA) along with their respective E3 treatment medium, with one larva per well and 24 larvae per treatment group. The plates were placed in lightproof boxes inside the incubator for overnight dark habituation. At 7 dpf, they were transferred to the Zebrabox (Viewpoint Life Sciences, Montréal, QC, Canada) to measure the visual-motor response (VMR) displayed by the larvae. The stimulating light intensity was 0.005 µW cm^−2^, or 0.01 lux (1.80e−5 µW cm−2 at 500 nm), to trigger scotopic VMR.^45^ The larval movement was recorded in tracking mode, which binned the activity every second. The data collection protocol consists of a 30-minute dark period, a 1-hour light period, and a 5-minute dark period. All VMR experiments were conducted between 9 a.m. and 6 p.m. to minimize the effect of circadian rhythm on vision. ^45,87^

### Enucleation

Enucleation was conducted at 5 dpf, following our established protocol.^43^ Before enucleation, larvae were anesthetized with 0.4 mg/mL Tricaine in Ringer’s solution ^88^ and placed on 1% agarose gel. The larval eyes were enucleated with a sterilized tungsten wire loop. The enucleated larvae were transferred to Ringer’s solution for a 2-hour recovery period prior to further drug treatment.

### Histology

The larvae used for histology were fixed in 4% paraformaldehyde (PFA) overnight at room temperature. The fixed larvae were sequentially infiltrated with 5%, 10%, 15%, and 20% sucrose for 1 hour each at room temperature, followed by overnight infiltration with 30% sucrose. These larvae were then embedded in the Tissue Freezing Medium (General Data, Cincinnati, OH). Ten-micrometer-thick cryosections were collected using a Leica CM1510S cryostat (Leica Biosystems, Deer Park, IL) and mounted onto Superfrost Plus Microscope Slides (Thermo Fisher Scientific, Rockford, IL). Rods were labeled by the *Tg(-3.7rho:EGFP)* transgene.^58^ Rod death was quantified using TUNEL staining, performed with the In Situ Cell Death Detection Kit, TMR red (Roche, Indianapolis, IN). Rod neogenesis was quantified by comparing *Tg(-3.7rho:EGFP)* larvae treated by hit compounds to those treated by 1.2 µM RA (MedChemExpress, Monmouth Junction, NJ). All sections were counterstained with 1 µg/mL DAPI solution (Thermo Fisher Scientific, Rockford, IL) and mounted using an anti-fade mounting medium (0.2% N-propyl gallate in 90% glycerol/1X PBS). The mounted sections were imaged using an Olympus BX51 microscope (Central Valley, PA) and a SPOT RT3 Color Slider camera (Sterling Heights, MI). The resulting images were processed by FIJI <https://imagej.net/software/fiji/>.

### Tapping Assay

A tapping assay was conducted to evaluate the total distance traveled by larvae in response to a mechanosensory stimulus, i.e., a physical tap. These larvae were transferred to 96-well plates and placed in the Zebrabox as described in the VMR assay section. To induce mechanosensory stimulation, a metal rod was manually tapped against the side of the 96-well plate under constant ambient light. Each experiment consists of 12 taps, with a 10-second interval between taps. Larval responses were recorded by the Zebrabox in tracking mode. Each plate was tested twice. During the recording, the larvae were exposed to constant ambient light.

## Data Analysis

### Data and Code Availability

The experimental data in this study can be accessed via the Harvard Dataverse at <https://dataverse.harvard.edu/dataverse/zebrafish_Q344X_FDA/>. The analysis code can be accessed through GitHub at <https://github.com/wang4537/Q344X_zebrafish_VMR_FDA>. All data analyses were conducted using R version 4.3.1 <https://www.r-project.org/>.

### Hit Identification from the VMR Screening Data

The VMR data were first normalized to remove confounding factors, including light-intensity variation across the 96-plate, batch effects across biological replications, and baseline variation across different sample types, using our established procedure. ^89^ Two approaches were then analyzed on the normalized data. In the first approach, clustering analysis was conducted using the average total distance for each sample type during the 5-minute dark period, from 0 to 29 seconds. Three clustering algorithms were used: HC, K-Means, and GMM-EM. In the clustering result, if the drug-treated Q344X groups clustered with the WT control groups, the corresponding drugs were classified as Type I hits. In HC and K-Means, the number of clusters was affected by linkage type and k values, respectively. For HC, both average and complete linkages were used to generate the clustering results, and the intersection of these results was selected as the set of hits identified by HC. For K-Means, k = 10, 20, and 30 were used to generate clustering results, and the intersection of these results was selected as the set of hits identified by K-Means. For GMM-EM, the optimal number of clusters was determined by the EM algorithm, and only one clustering result was generated to identify hits. In the second approach, the Welch two-sample t-test was used to compare the differences of average total distance traveled by drug-treated and control Q344X larvae at the first second of the 5-minute dark period. If the drug-treated Q344X larvae exhibited a significant increase in average total distance traveled compared with the DMSO-treated ones (Bonferroni-adjusted p-value < 0.05), the corresponding drug was classified as a Type II hit.

### Analysis of Other VMR Experiments

For the other VMR experiments, the data were normalized as in the VMR screening analysis above. Then, the pairwise Wilcoxon rank-sum test was used to compare the average total distance traveled by larvae in the first second seconds after light offset between the two treatment groups. The resulting p-values were adjusted using the Bonferroni method to correct for multiple comparisons.

### Histological Analysis

Cells with positive EGFP or TUNEL signals were counted from both retinal sections of each larva and averaged. Positive cell counts between treatment groups were compared using the Wilcoxon rank-sum test. The resulting p-values were adjusted using Holm’s method to correct for multiple comparisons.

### Tapping Assay Analysis

The tapping assay data were analyzed in two steps: 1) filtering and 2) hypothesis testing. First, the total distance data were filtered to remove unsuccessful taps. A tap was considered unsuccessful if the average total distance traveled at that tap was less than two standard deviations above the average total distance traveled during periods between taps. The average total distance during the in-between periods was calculated across all treatment groups and all in-between time intervals. Second, the filtered data from all taps and plates were pooled within each treatment group. The Wilcoxon rank-sum test was used to test the difference in average total distance traveled between groups. The resulting p-values were adjusted using Holm’s method to correct for multiple comparisons.^90^

### Chemical Structure Similarity Analysis

The structural similarities between hits were analyzed based on their Simplified Molecular-Input Line Entry System (SMILES) representations (provided by the vendor, SelleckChem). Using the hit SMILES, the chemical structure fingerprints were extracted using the *rcdk* package (version 3.8.1) from R,^91^ and structural similarities were analyzed using HC.

## ACKNOWLEDGEMENT

We acknowledge the use of the Chemical Genomics Facility (CGF), a core facility of Purdue Institute for Drug Discovery and the NIH-funded Indiana Clinical and Translational Sciences Institute. In particular, we thank Dr. Li Wu and Dr. Lan Chen for their support on instrumentations and the compound library. This publication was made possible, in part, with support from the Indiana Clinical and Translational Sciences Institute to YFL and BW, funded in part by Grant Number UM1TR004402 from the National Institutes of Health, National Center for Advancing Translational Sciences, Clinical and Translational Sciences Award. LG was supported by Grant Numbers TL1 TR001107 and UL1 TR001108 (A. Shekhar, PI) from the National Institutes of Health, National Center for Advancing Translational Sciences, Clinical and Translational Sciences Award. The content is solely the responsibility of the authors and does not necessarily represent the official views of the National Institutes of Health. BW, LG, and YFL were partially supported by research grants from Purdue Institute for Drug Discovery. MT was supported by Japan Society for the Promotion of Science grants JSPS KAKENHI 23K21480 and JSPS KAKENHI 24K22167. YFL was partially supported by grants from the Purdue Research Foundation and the International Retinal Research Foundation. The graphic abstract was created in BioRender. Wang, B. (2026) https://BioRender.com/hkw78kb.

## AUTHOR CONTRIBUTIONS

**B.W.:** Conceptualization, Methodology, Software, Validation, Formal Analysis, Investigation, Data Curation, Visualization, Writing – original draft, Writing – review & editing, Visualization. **L.G.:** Conceptualization, Methodology, Software, Investigation, Data Curation, Writing – review & editing, Methodology, Conceptualization, Funding Acquisition. **E.C.:** Investigation, Writing – review & editing, Methodology, Conceptualization. **R.J.:** Investigation. **T.K.:** Investigation. **X.Z.:** Investigation. **J.A.N.:** Investigation. **M.T.:** Methodology, Resources. **Y.F.L.:** Conceptualization, Resources, Writing – original draft, Writing – review & editing, Supervision, Project Administration, Funding acquisition.

