## Supplemental Figures and Tables for "Zebrafish Functional Screening of FDA-Approved Drugs for Autosomal Dominant Retinitis Pigmentosa Caused by RHODOPSIN Q344X Mutation" (from another version of this preprint: version 1, 2026-04-21)

**Supplemental Information**

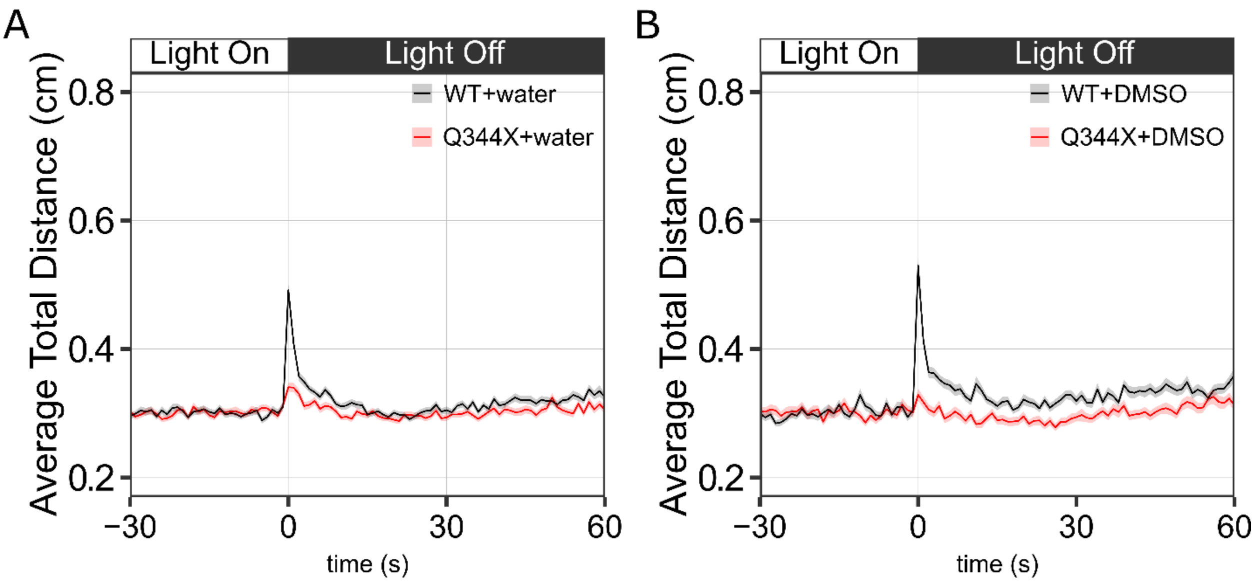


**Supp Figure 1. Average total distance curves of WT and Q344X larvae in the light-off VMR assay.**

**(A)** Average total distance curves of water-treated WT and Q344X larvae during light offset. The x-axis of the plot indicates the time window, spanning from 30 seconds before light offset (-30 s) to 60 seconds following it (60 s). Sample size: 48 larvae × 18 biological replicates per genotype. Q344X larvae (red) exhibited a significant reduction in average total distance traveled (y-axis) compared with WT larvae (black) during the first second following light offset (Welch two-sample test, test statistics = -13.2983, df = 1601,7, p-value = 2.36e-38). **(B)** Average total distance curves of DMSO-treated WT and Q344X larvae. The x-axis of the plot indicates the time window, spanning from 30 seconds before light offset (-30 s) to 60 seconds following it (60 s). Sample size: 48 larvae × 9 biological replicates per genotype. Q344X larvae (red) exhibited a significant reduction in average total distance traveled compared with WT larvae (black) during the first second following light offset (Welch two-sample test, test statistics = -12.4022, df =

77.452, p-value = 2.34e-32). In both plots, data were normalized using established methods.[83] The ribbons indicate the standard error of the mean. The white and black bars above the plots denote the light-on and light-off phases of the time course, respectively.

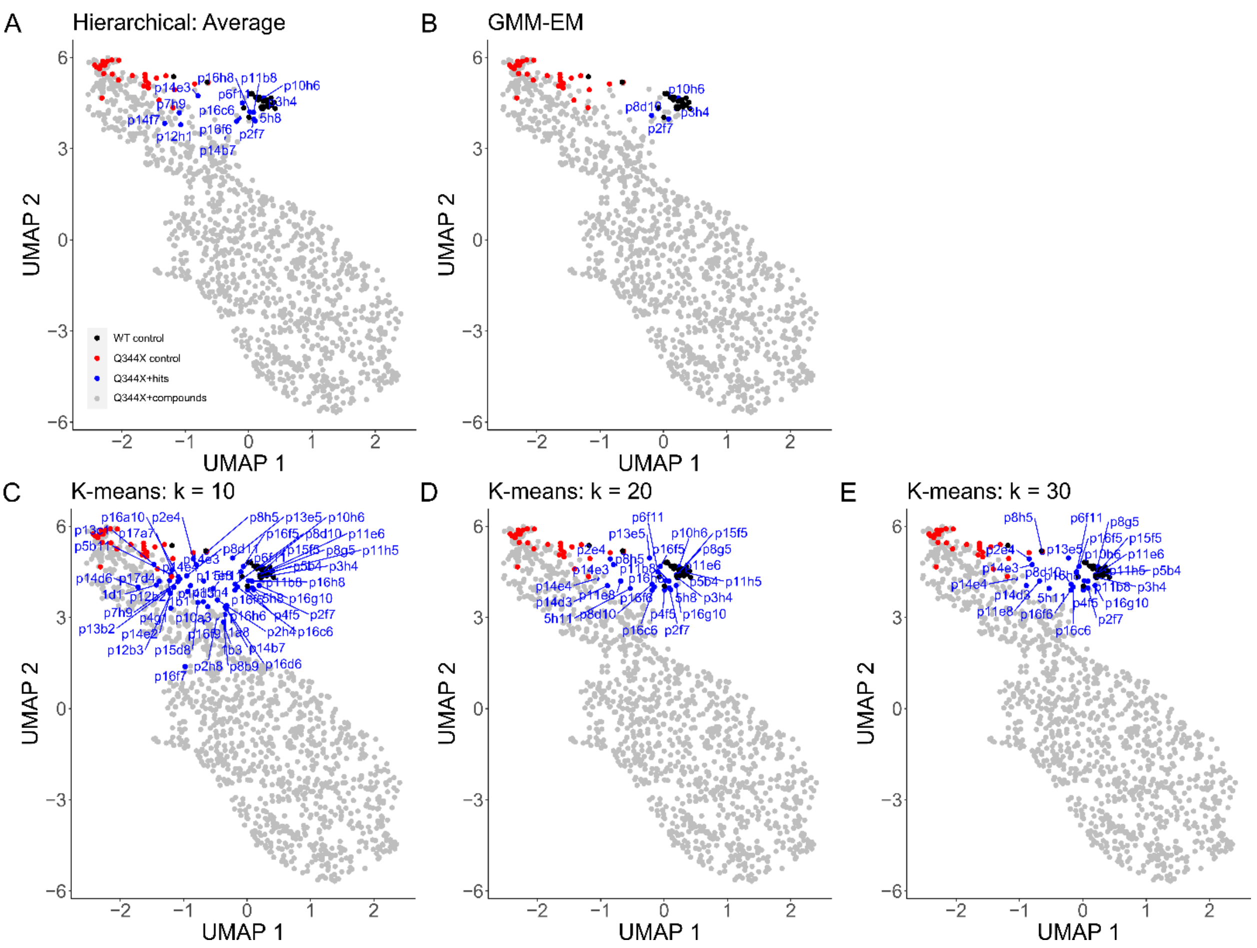


**Supp Figure 2. UMAP plots of additional clustering algorithms for the identification of Type I hits.**

**(A)** Identification of Type I hits using hierarchical clustering with average linkage. (B) Identification of Type I hits using a Gaussian Mixture Model with Expectation Maximization (GMM-EM) (C–E) Identification of Type I hits using k-means clustering using k = 10 (C), k = 20 (D), and k = 30 (E). Four groups are colored: WT controls (black), Q344X controls (red), drug-treated Q344X groups clustered with WT controls based on the corresponding clustering algorithm (Q344X+hits; blue), and the remaining drug-treated Q344X groups (Q344X+compouns; gray). The codenames of the Type I hits are labeled next to the Q344X groups treated with those compounds. The clustering results across different hyperparameters within one algorithm were intersected to yield the hits identified by each algorithm.

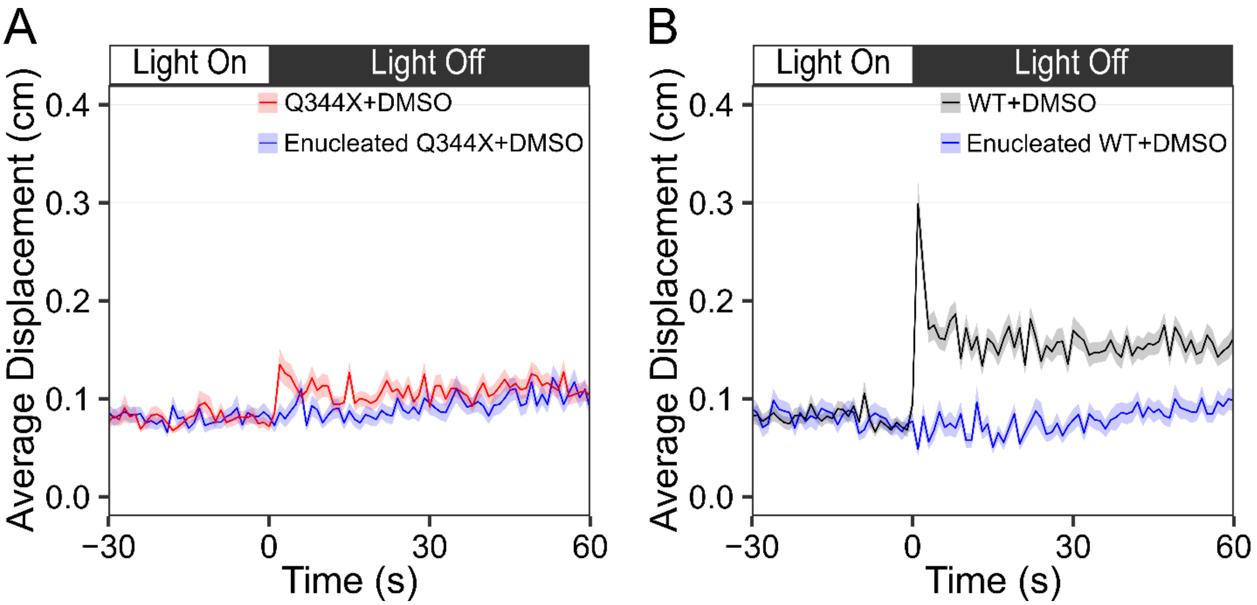


**Supp Figure 3. Average total distance curves of WT and Q344X control larvae, with or without enucleation, in the light-off VMR assay.**

**(A)** Averaged total distance curves of DMSO-treated Q344X larvae with (blue) and without enucleation (red). DMSO-treated Q344X larvae without enucleation (red) exhibited a significant increase in average total distance traveled compared with DMSO-treated Q344X larvae with enucleation (blue), during the first second following light offset (Wilcoxon rank-sum test result: test statistic = 9070, p-value = 1.49e-08, adj. p-value = 2.98e-08). **(B)** Averaged total distance curves of DMSO-treated WT larvae with (blue) and without enucleation (black). DMSO-treated WT larvae without enucleation (black) exhibited a significant increase in average total distance traveled, compared with DMSO-treated WT larvae with enucleation (blue), during the first second following light offset (Wilcoxon rank-sum test result: test statistic = 2398, p-value = 1.55e-39, adj. p-value = 3.1e-39). Sample size: 24 larvae × 6 biological replicates for each of the following groups—Q344X+DMSO, enucleated Q344X+DMSO, WT+DMSO, and enucleated WT+DMSO. In these plots, the ribbons indicate the standard error of the mean. The white and black bars above the plots denote the light-on and light-off phases of the time course, respectively.

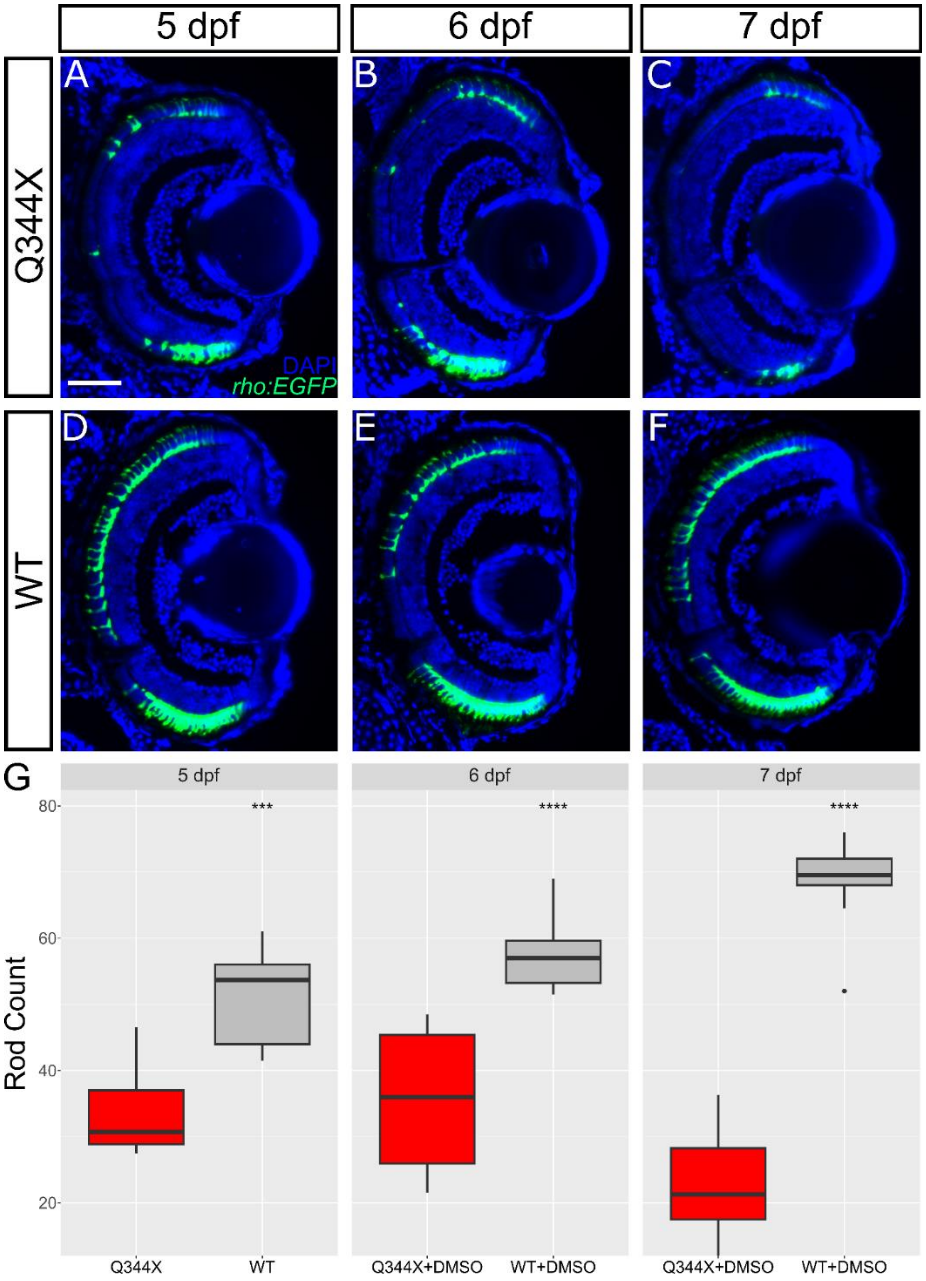


**Supp Figure 4. Rods in WT and Q344X retinas at 5, 6, and 7 dpf under control conditions.**

**(A–C)** Representative images of retinal cryosections from Q344X larvae at 5, 6, and 7 dpf, respectively. DMSO was applied at 10 μM from 5 to 7 dpf, consistent with the VMR screening protocol. Samples at 5 dpf were collected prior to DMSO treatment. Rods were visualized using the *Tg(-3.7rho:EGFP)* transgene. Nuclei were counterstained with DAPI. Scale bar: 50 µm. **(D–F)** Representative images of retinal cryosections of WT larvae at 5, 6, and 7 dpf, respectively. Samples were processed as described in the Q344X samples above. (G) Box-and-whisker plots showing rod counts at 5, 6, and 7 dpf in Q344X (red) and WT (gray) retinas. Statistical test results are shown in Supp Table 3. Significance levels: ***, p-value < 0.001; ****, p-value < 0.0001.

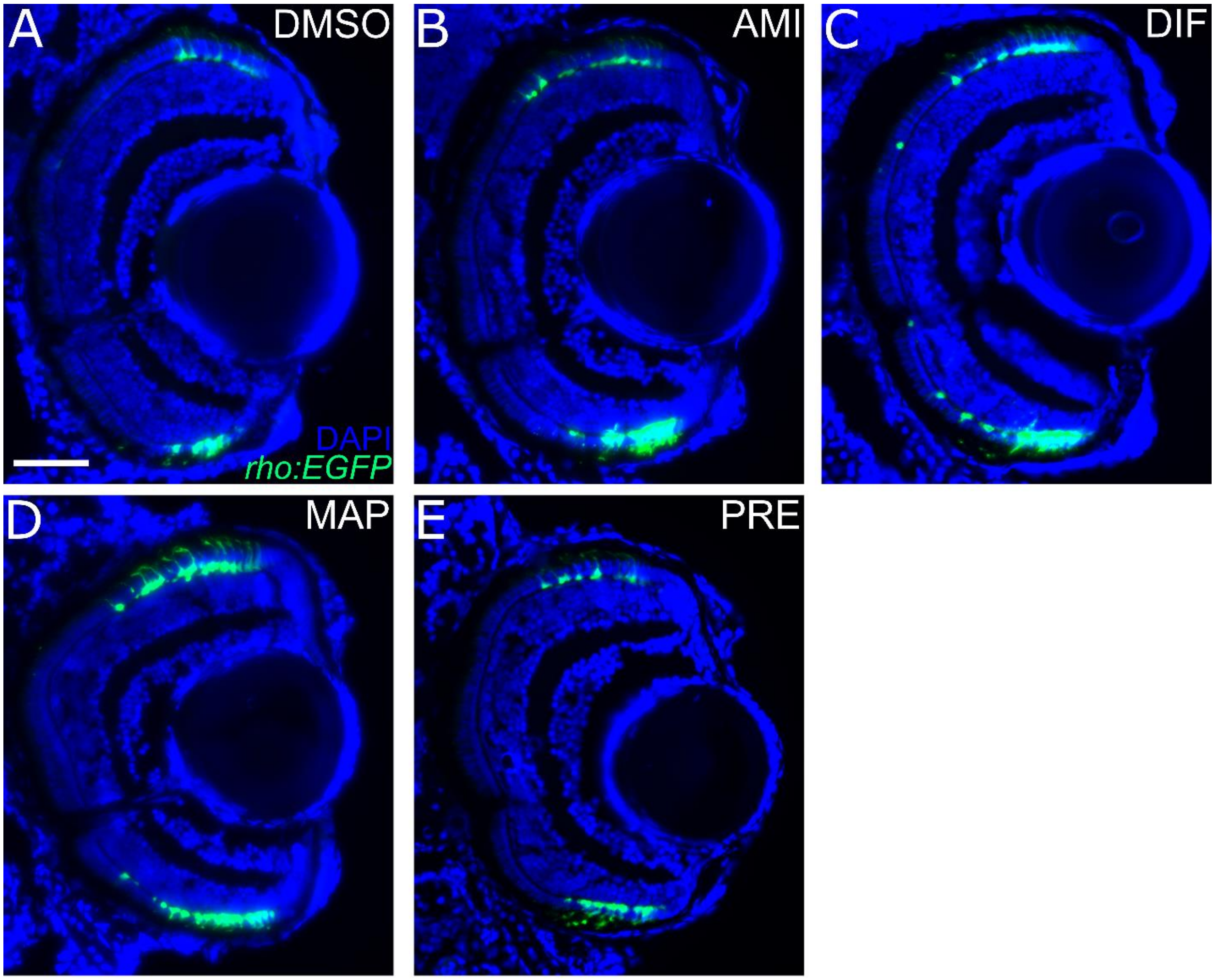


**Supp Figure 5. Rods in Q344X retinas at 7 dpf after hit treatment.**

**(A–E)** Representative images of retinal cryosections from 7-dpf Q344X larvae following treatment with DMSO (A), AMI (B), DIF (C), MAP (D), and PRE (E). Treatments were applied at 10 μM from 5 to 7 dpf, consistent with the VMR screening protocol. Rods were visualized using the *Tg(-3.7rho:EGFP)* transgene. Nuclei were counterstained with DAPI. Scale bar: 50 μm. The corresponding quantification results can be found in **Error! Reference source not found.**A. Refer to Supp Figure 4C for the representative image of Q344X larvae treated by DMSO.

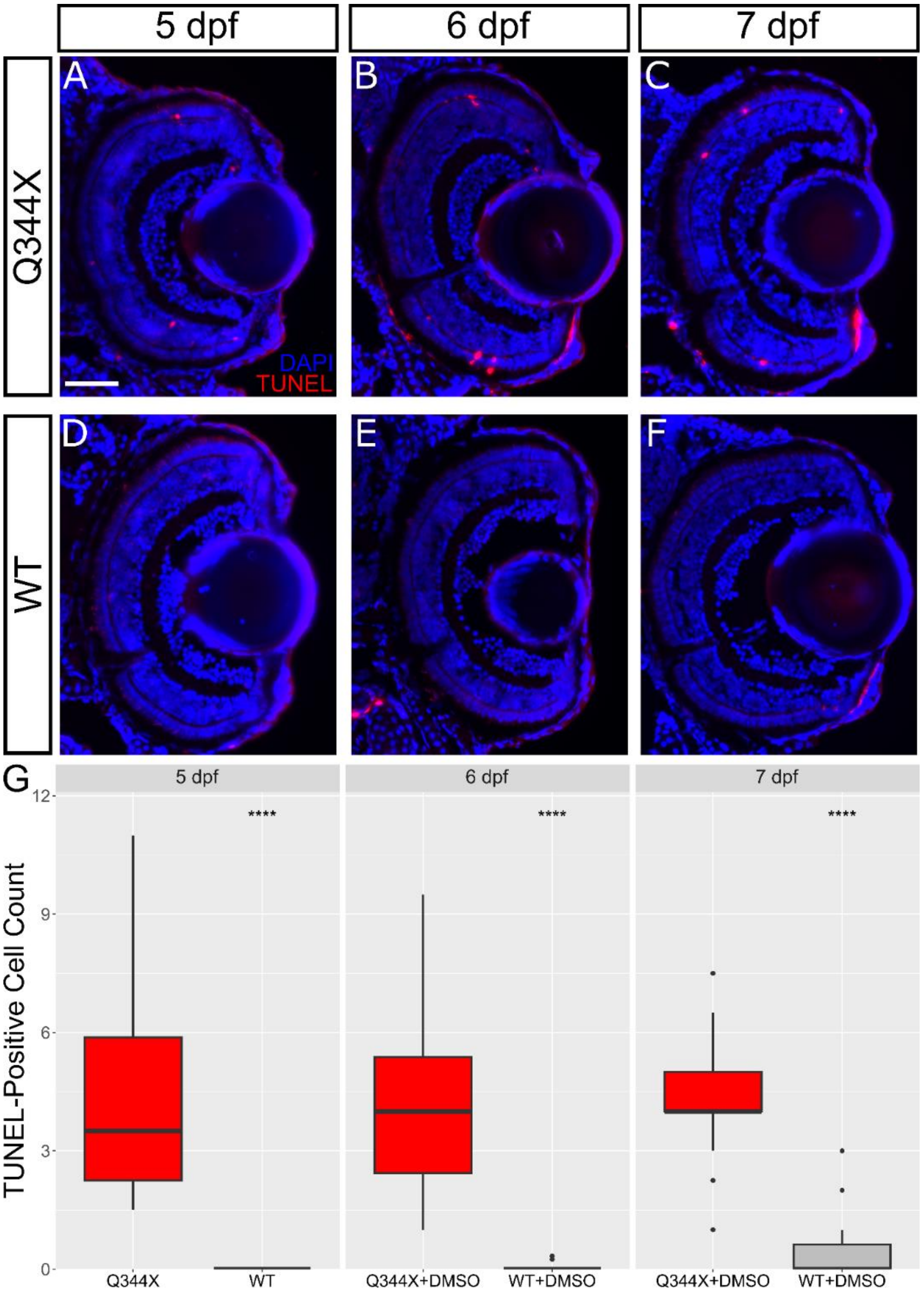


**Supp Figure 6. TUNEL-positive cells in WT and Q344X retinas at 5, 6, and 7 dpf under control condition.**

**(A–C)** Representative images of retinal cryosections from Q344X larvae at 5, 6, and 7 dpf, respectively. DMSO was applied at 10 μM from 5 to 7 dpf, consistent with the VMR screening protocol. Samples at 5 dpf were collected prior DMSO treatment. TUNEL-positive cells were visualized on cryosections according to manufacturer's protocols. Nuclei were counterstained with DAPI. Scale bar: 50 µm. **(D–F)** Representative images of retinal cryosections from WT larvae at 5, 6, and 7 dpf, respectively. Samples were processed as described in Q344X samples above. **(G)** Box-and-whisker plots showing TUNEL-positive cell counts at 5, 6, and 7 dpf in Q344X (red) and WT (gray) retinas. Statistical test results are shown in Supp Table 5. Significance level: ****, p-value < 0.0001.

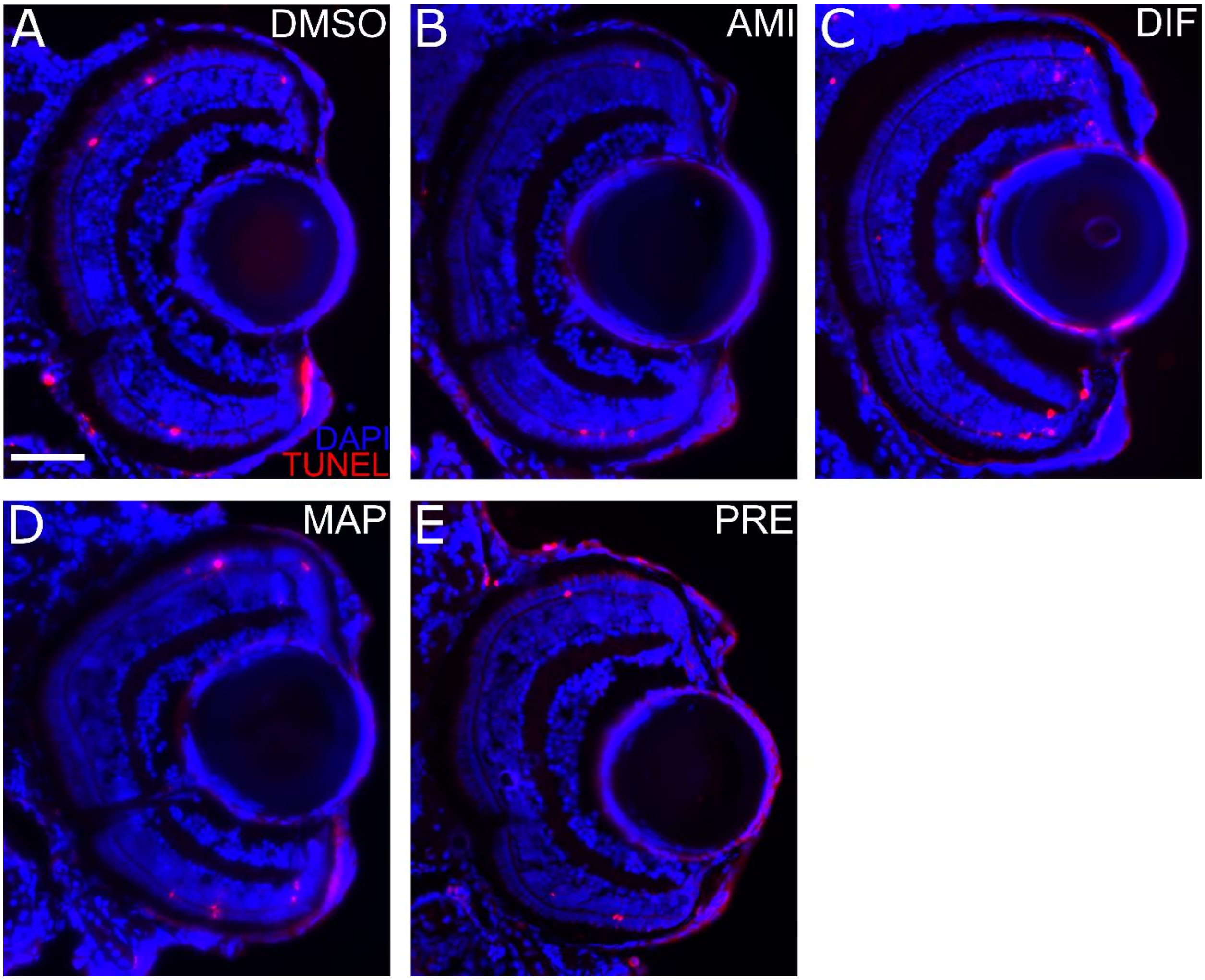


**Supp Figure 7. TUNEL-positive cells in WT and Q344X retinas at 7 dpf after hit treatment.**

**(A–E)** Representative images of TUNEL staining on retinal cryosections from 7-dpf Q344X larvae following treatment with DMSO (A), AMI (B), DIF (C), MAP (D), and PRE (E). Treatments were applied at 10 μM from 5 to 7 dpf, consistent with the VMR screening protocol. TUNEL-positive cells were visualized on cryosections according to the manufacturer's protocols. The nuclei were counterstained with DAPI. Scale bar: 50 μm. The corresponding quantification results can be found in **Error! Reference source not**

**found.**B. Refer to Supp Figure 6C for the representative image of Q344X larvae treated by DMSO.

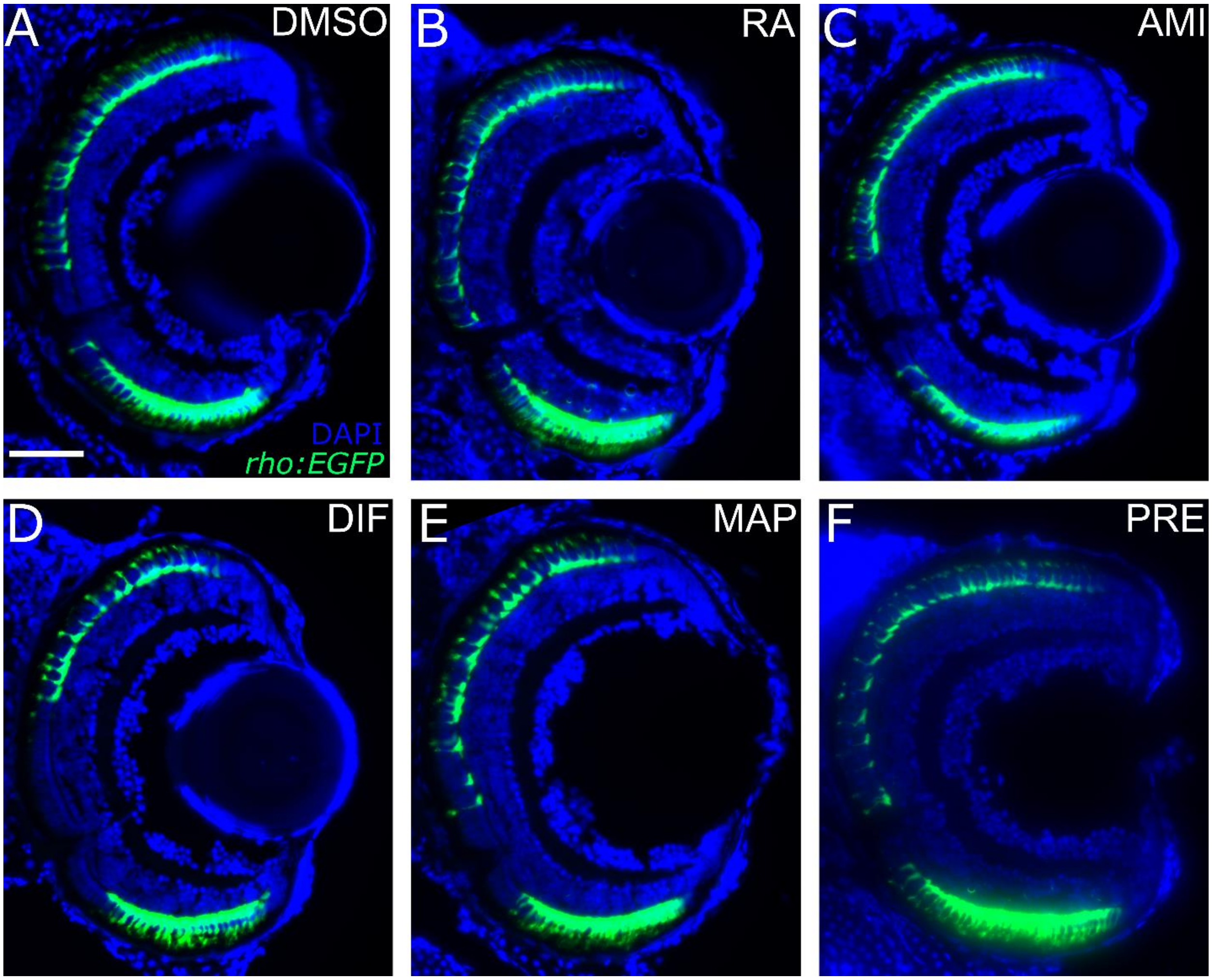


**Supp Figure 8. Rods in WT retinas at 7 dpf after retinoic acid (RA) and hit treatment.**

**(A–E)** Representative images of retinal cryosections of 7-dpf WT larvae following treatment with DMSO (A), RA (B), AMI (C), DIF (D), MAP (E), and PRE (F). Hit compounds were applied at 10 μM, and RA at 1.2 μM. All treatments were administered from 5 to 7 dpf, consistent with the VMR screening protocol. Rods were visualized using the transgene *Tg(-3.7rho:EGFP)*. The nuclei were counterstained with DAPI. Scale bar:

50 μm. The corresponding quantification results can be found in **Error! Reference source not found.**C.

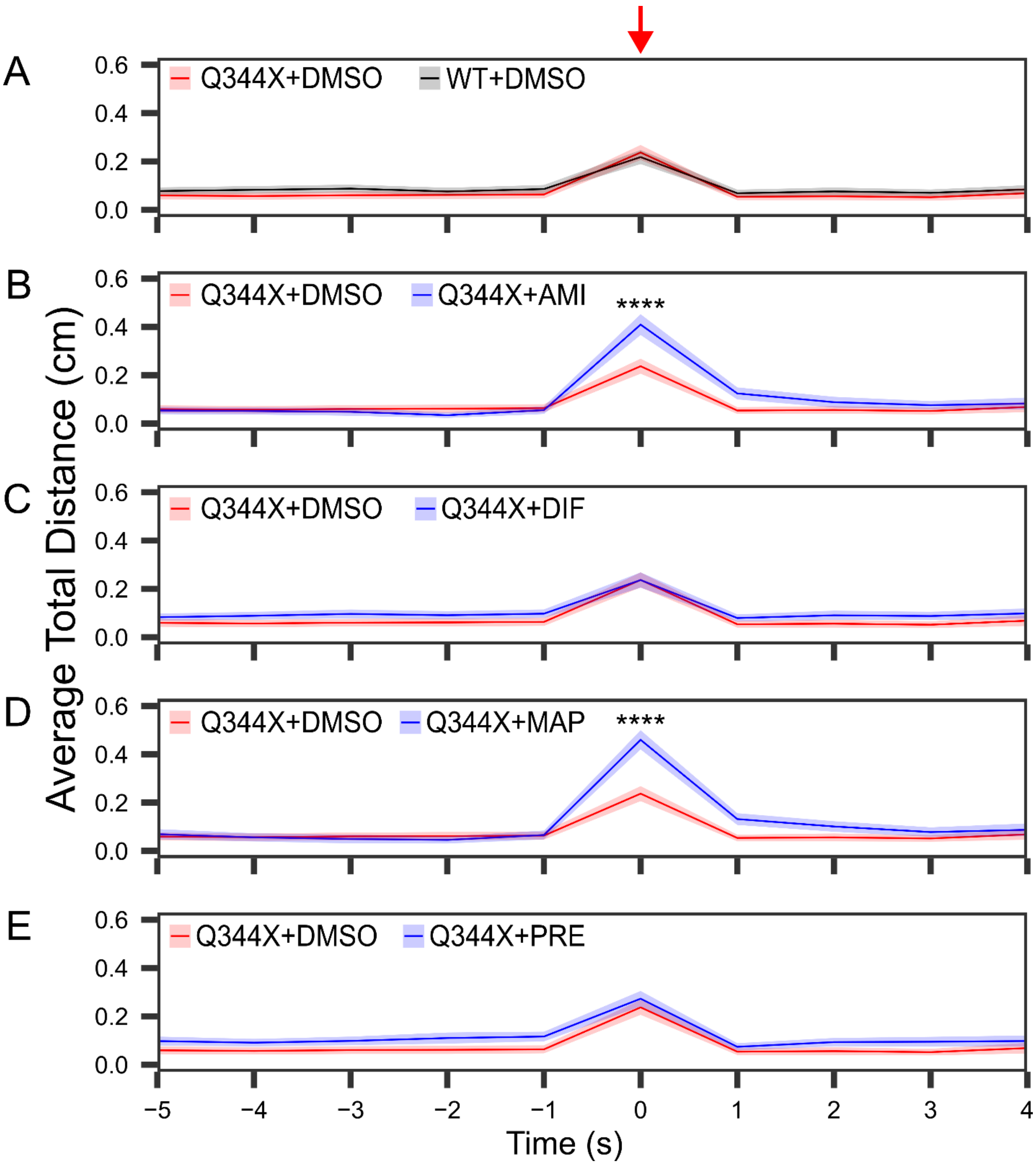


**Supp Figure 9. Average total distance curves of larvae treated by hit compounds in response to mechanosensory stimulation.**

**(A–E)** DMSO-treated Q344X larvae compared with DMSO-treated WT larvae (A), and Q344X larvae treated with AMI (B), DIF (C), MAP (D), or PRE(E). Each total distance curve represents the average total distance across all larvae per group, aggregated over 11 taps per repeat, 2 repeats per biological replicate, and 4 biological replicates in total. The red arrow indicates the time point at which the tap occurred. Unsuccessful trials were removed during data analysis (See Methods for details). In these plots, the ribbons indicate the standard error of the mean. Statistical results are shown in Supp Table 9. Significant level: ****, p-value $< 0.0001$.

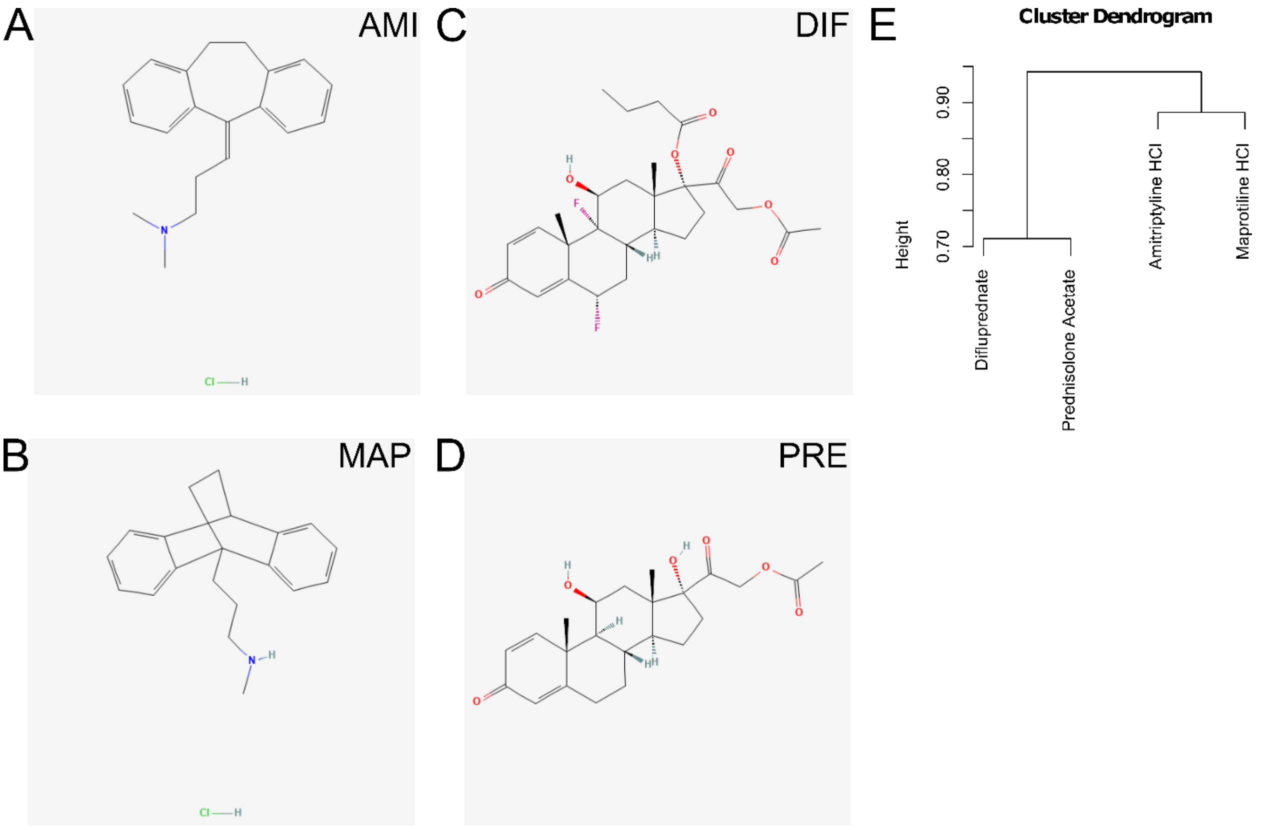


**Supp Figure 10. Chemical structures of the hit compounds and their structural similarity.**

**(A–D)** Chemical structures of AMI (A), MAP (B), DIF (C), and PRE (D). Structural information was retrieved from Pubchem (PubChem Identifies: AMI, CID 11065; MAP, CID 71478; DIF, CID 443936; PRE, CID 5834). **(E)** Dendrogram generated by hierarchical clustering using the chemical fingerprints of the hit compounds. These chemical fingerprints were extracted from SMILE data provided in the information sheet from the library vendor (SelleckChem). Hierarchical clustering was performed using complete linkage to construct the tree.

**Supp Table 1. List of compounds and vendors used in confirmatory testing and characterization.**

| Name | Code name | Vendor |
|---|---|---|
| Amitriptyline HCl | p10h6 | MedChemExpress, Sigma Millipore |
| Avagacestat (BMS-708163) | p2e4 | MedChemExpress |
| Bephenium Hydroxynaphthoate | p13e5 | MedChemExpress |
| Cephalothin | p14e3 | MedChemExpress |
| Chlormadinone acetate | p14d3 | MedChemExpress |
| Chlorprothixene | p5b4 | MedChemExpress, Sigma Millipore |
| Cilomilast | p3h4 | MedChemExpress |
| Desvenlafaxine Succinate | p11b8 | MedChemExpress |
| Difluprednate | p11e6 | MedChemExpress |
| Ellagic acid | p2f7 | MedChemExpress, SelleckChem |
| Etidronate | p16h8 | MedChemExpress |
| Gabapentin HCl | p16g10 | MedChemExpress |
| Gallic acid | p14e4 | MedChemExpress |
| Histamine 2HCl | p11e8 | MedChemExpress |
| Loxapine Succinate | p11h5 | MedChemExpress |
| Maprotiline HCl | p8h5 | MedChemExpress |
| Oxytocin (Syntocinon) | p16f5 | MedChemExpress |
| Pamidronate Disodium | p16f6 | MedChemExpress |
| Pemetrexed | p16c6 | MedChemExpress |
| Phenylbutazone | p4f5 | MedChemExpress |
| Potassium Iodide | p5h11 | Flinn Chemicals |
| Tetracycline HCl | p8d10 | MedChemExpress |
| Tiopronin | p6f11 | MedChemExpress |
| Vidofludimus | p15f5 | MedChemExpress |
| Xylazine HCl | p8g5 | MedChemExpress |

**Supp Table 2. Pairwise comparisons of the average total distance of Q344X larvae with or without enucleation, following hit or DMSO treatment, during the first second following light offset in the VMR assay.**

Statistical comparisons were performed using the Welch's two-sample t-test, and multiple comparisons were adjusted using the Bonferroni method. The corresponding VMR results are shown in Error! Reference source not found..

Significance levels: ns, not significant; *, p-value < 0.05; **, p-value < 0.01.

| Hit | Group1 | Mean1 | SD1 | n1 | Group2 | Mean2 | SD2 | n2 | statistic | df | p | p.adj | significance |
|---|---|---|---|---|---|---|---|---|---|---|---|---|---|
| AMI | Q344X+AMI | 0.145 | 0.2 | 72 | Q344X+DMSO | 0.081 | 0.089 | 168 | 2.614923 | 83.39072 | 0.011 | 0.033 | * |
| | Q344X_Enu+AMI | 0.081 | 0.069 | 72 | Q344X+AMI | 0.145 | 0.2 | 72 | -2.55106 | 87.59445 | 0.013 | 0.039 | * |
| | Q344X_Enu+AMI | 0.081 | 0.069 | 72 | Q344X_Enu+DMSO | 0.077 | 0.095 | 168 | 0.398779 | 183.072 | 0.691 | 1 | ns |
| DIF | Q344X+DIF | 0.197 | 0.289 | 48 | Q344X+DMSO | 0.081 | 0.089 | 168 | 2.741453 | 49.59329 | 0.008 | 0.024 | * |
| | Q344X_Enu+DIF | 0.045 | 0.097 | 48 | Q344X+DIF | 0.197 | 0.289 | 48 | -3.45258 | 57.37762 | 0.001 | 0.003 | ** |
| | Q344X_Enu+DIF | 0.045 | 0.097 | 48 | Q344X_Enu+DMSO | 0.077 | 0.095 | 168 | -2.04173 | 75.19864 | 0.045 | 0.135 | ns |
| MAP | Q344X+MAP | 0.201 | 0.265 | 48 | Q344X+DMSO | 0.081 | 0.089 | 168 | 3.090269 | 50.08029 | 0.003 | 0.009 | ** |
| | Q344X_Enu+MAP | 0.087 | 0.108 | 48 | Q344X+MAP | 0.201 | 0.265 | 48 | -2.74834 | 62.02984 | 0.008 | 0.024 | * |
| | Q344X_Enu+MAP | 0.087 | 0.108 | 48 | Q344X_Enu+DMSO | 0.077 | 0.095 | 168 | 0.6074 | 69.53015 | 0.546 | 1 | ns |
| PRE | Q344X+PRE | 0.237 | 0.284 | 48 | Q344X+DMSO | 0.081 | 0.089 | 168 | 3.755367 | 49.68599 | 0.000455 | 0.001365 | ** |
| | Q344X_Enu+PRE | 0.082 | 0.119 | 48 | Q344X+PRE | 0.237 | 0.284 | 48 | -3.48461 | 63.04733 | 0.000902 | 0.002706 | ** |
| | Q344X_Enu+PRE | 0.082 | 0.119 | 48 | Q344X_Enu+DMSO | 0.077 | 0.095 | 168 | 0.264585 | 65.18579 | 0.792 | 1 | ns |

**Supp Table 3. Pairwise comparisons of rod counts at 5, 6, and 7 dpf between Q344X and WT retinas.**

Statistical comparisons were performed using the Wilcoxon rank-sum test, and multiple comparisons were corrected using Holm's method. The corresponding images and plots of these count data are presented in Supp Figure 4. Significance levels: ***, p-value < 0.001; ****, p-value < 0.0001.

| Stage | Group1 | Mean1 | SD1 | n1 | Group2 | Mean2 | SD2 | n2 | statistic | p | p.adj | significance |
|---|---|---|---|---|---|---|---|---|---|---|---|---|
| 5 dpf | Q344X | 33.3383 | 6.3734 | 10 | WT | 51.5128 | 7.0345 | 10 | 4 | 0.000175 | 0.000175 | *** |
| 6 dpf | Q344X+DMSO | 35.0365 | 9.9906 | 16 | WT+DMSO | 57.7652 | 6.0952 | 16 | 0 | 1.56E-05 | 4.68E-05 | **** |
| 7 dpf | Q344X+DMSO | 22.6190 | 7.1505 | 14 | WT+DMSO | 68.6212 | 6.3399 | 14 | 0 | 2.76E-05 | 5.52E-05 | **** |

**Supp Table 4. Pairwise comparisons of rod counts at 7 dpf between DMSO- and hit-treated Q344X retinas.**

Statistical comparisons were performed using the Wilcoxon rank-sum test, and multiple comparisons were corrected using Holm's method. The corresponding images and count data are presented in **Error! Reference source not found.**.

Significance levels: ns, not significant; **, p-value < 0.01.

| Group1 | Mean1 | SD1 | n1 | Group2 | Mean2 | SD2 | n2 | statistic | p | p.adj | significance |
|---|---|---|---|---|---|---|---|---|---|---|---|
| Q344X+DMSO | 22.6190 | 7.1505 | 14 | Q344X+AMI | 29.0278 | 6.9346 | 18 | 68 | 0.028 | 0.084 | ns |
| Q344X+DMSO | 22.6190 | 7.1505 | 14 | Q344X+DIF | 37.4111 | 9.6226 | 15 | 25.5 | 0.000559 | 0.002236 | ** |
| Q344X+DMSO | 22.6190 | 7.1505 | 14 | Q344X+MAP | 25.7353 | 7.4638 | 17 | 98.5 | 0.427 | 0.854 | ns |
| Q344X+DMSO | 22.6190 | 7.1505 | 14 | Q344X+PRE | 23.0758 | 3.5072 | 11 | 73 | 0.848 | 0.854 | ns |

**Supp Table 5. Pairwise comparisons of TUNEL-positive cell counts at 5, 6, and 7 dpf between Q344X and WT retinas.**

Statistical comparisons were performed using the Wilcoxon rank-sum test, and multiple comparisons were corrected using Holm's method. The corresponding images and count data are presented in Supp Figure 6. Significance level: ****, p-value < 0.0001.

| Stage | Group1 | Mean1 | SD1 | n1 | Group2 | Mean2 | SD2 | n2 | statistic | p | p.adj | significance |
|---|---|---|---|---|---|---|---|---|---|---|---|---|
| 5 dpf | Q344X | 4.4 | 2.865504 | 10 | WT | 0 | 0 | 13 | 130 | 9.83E-06 | 9.83E-06 | **** |
| 6 dpf | Q344X+DMSO | 4.022222 | 2.226832 | 30 | WT+DMSO | 0.05303 | 0.119447 | 11 | 330 | 1.08E-06 | 3.24E-06 | **** |
| 7 dpf | Q344X+DMSO | 4.464286 | 1.553797 | 21 | WT+DMSO | 0.541667 | 0.987613 | 12 | 247.5 | 4.50E-06 | 9.00E-06 | **** |

**Supp Table 6. Pairwise comparisons of TUNEL-positive cell counts at 7 dpf between DMSO-treated controls and hit-treated Q344X retinas in the outer nuclear layer.**

Statistical comparisons were performed using the Wilcoxon rank-sum test, and multiple comparisons were adjusted using Holm's method. The corresponding images and count data are presented in **Error! Reference source not found.**B.

Significance levels: ns, not significant; ***, p-value < 0.001.

| Group1 | Mean1 | SD1 | n1 | Group2 | Mean2 | SD2 | n2 | statistic | p | p.adj | significance |
|---|---|---|---|---|---|---|---|---|---|---|---|
| Q344X+DMSO | 4.4643 | 1.5538 | 21 | Q344X+AMI | 3.6500 | 1.9518 | 13 | 182 | 0.109 | 0.207 | ns |
| Q344X+DMSO | 4.4643 | 1.5538 | 21 | Q344X+DIF | 3.5000 | 2.0303 | 11 | 159.5 | 0.082 | 0.207 | ns |
| Q344X+DMSO | 4.4643 | 1.5538 | 21 | Q344X+MAP | 3.4015 | 1.4714 | 11 | 161 | 0.069 | 0.207 | ns |
| Q344X+DMSO | 4.4643 | 1.5538 | 21 | Q344X+PRE | 1.3788 | 0.9577 | 11 | 221.5 | 2.53E-05 | 0.000101 | *** |

**Supp Table 7. Pairwise comparisons of rod counts at 7 dpf between WT retinas treated with DMSO, RA, or hit compounds.**

Statistical comparisons were performed using the Wilcoxon rank-sum test, and multiple comparisons were adjusted using the Bonferroni method. The corresponding image and count data are presented in **Error! Reference source not found.**C. Significance levels: ns, not significant; *, p-value < 0.05; **, p-value < 0.01.

| Group1 | Mean1 | SD1 | n1 | Group2 | Mean2 | SD2 | n2 | statistic | p | p.adj | significance |
|---|---|---|---|---|---|---|---|---|---|---|---|
| WT+DMSO | 68.6212 | 6.3399 | 11 | WT+RA | 77.0444 | 8.3529 | 15 | 33.5 | 0.012 | 0.027 | * |
| WT+DMSO | 68.6212 | 6.3399 | 11 | WT+AMI | 75.7889 | 8.0063 | 15 | 27 | 0.004 | 0.016 | * |
| WT+DMSO | 68.6212 | 6.3399 | 11 | WT+MAP | 59.4559 | 6.6111 | 17 | 161 | 0.002 | 0.01 | ** |
| WT+DMSO | 68.6212 | 6.3399 | 11 | WT+DIF | 61.7794 | 6.4041 | 17 | 149.5 | 0.009 | 0.027 | * |
| WT+DMSO | 68.6212 | 6.3399 | 11 | WT+PRE | 67.8690 | 6.5653 | 14 | 83 | 0.763 | 0.763 | ns |

**Supp Table 8. Pairwise comparisons of average total distance in the WT with or without enucleation after hit or DMSO treatment during the first second after light offset in the VMR assay.**

Statistical comparisons were performed using the Welch's t test, and multiple comparisons were adjusted using the Bonferroni method. The corresponding data are presented in **Error! Reference source not found.**. Significance levels: ns, not significant; *, p-value < 0.05; **, p-value < 0.01; ****, p-value < 0.0001.

| Hit | Group1 | Mean1 | SD1 | n1 | Group2 | Mean2 | SD2 | n2 | statistic | df | p | p.adj | significance |
|---|---|---|---|---|---|---|---|---|---|---|---|---|---|
| AMI | WT+Ami | 0.199 | 0.26 | 72 | WT+DMSO | 0.3 | 0.292 | 168 | -2.65063 | 149.8444 | 0.009 | 0.027 | * |
| | WT-Enu+Ami | 0.089 | 0.175 | 72 | WT+Ami | 0.199 | 0.26 | 72 | -2.99223 | 124.4494 | 0.003 | 0.009 | ** |
| | WT-Enu+Ami | 0.089 | 0.175 | 72 | WT-Enu+DMSO | 0.052 | 0.099 | 168 | 1.669663 | 90.88254 | 0.098 | 0.294 | ns |
| DIF | WT+DIF | 0.33 | 0.331 | 48 | WT+DMSO | 0.3 | 0.292 | 168 | 0.566003 | 69.26096 | 0.573 | 1 | ns |
| | WT-Enu+DIF | 0.073 | 0.137 | 48 | WT+DIF | 0.33 | 0.331 | 48 | -4.97447 | 62.62541 | 5.39E-06 | 1.62E-05 | **** |
| | WT-Enu+DIF | 0.073 | 0.137 | 48 | WT-Enu+DMSO | 0.052 | 0.099 | 168 | 0.985533 | 61.5944 | 0.328 | 0.984 | ns |
| MAP | WT+MAP | 0.344 | 0.253 | 48 | WT+DMSO | 0.3 | 0.292 | 168 | 1.031264 | 86.0209 | 0.305 | 0.915 | ns |
| | WT-Enu+MAP | 0.101 | 0.192 | 48 | WT+MAP | 0.344 | 0.253 | 48 | -5.31017 | 87.71083 | 8.18E-07 | 2.45E-06 | **** |
| | WT-Enu+MAP | 0.101 | 0.192 | 48 | WT-Enu+DMSO | 0.052 | 0.099 | 168 | 1.691765 | 54.2389 | 0.096 | 0.288 | ns |
| PRE | WT+PRE | 0.14 | 0.196 | 48 | WT+DMSO | 0.3 | 0.292 | 168 | -4.42233 | 112.5548 | 2.26E-05 | 6.78E-05 | **** |
| | WT-Enu+PRE | 0.043 | 0.132 | 48 | WT+PRE | 0.14 | 0.196 | 48 | -2.83515 | 82.17039 | 0.006 | 0.018 | * |
| | WT-Enu+PRE | 0.043 | 0.132 | 48 | WT-Enu+DMSO | 0.052 | 0.099 | 168 | -0.42205 | 62.83054 | 0.674 | 1 | ns |

**Supp Table 9. Pairwise comparison of average total distance at the first second after the tap stimulation in control and hit-treated larvae.**

The hypothesis testing used the Wilcoxon rank-sum test, and multiple hypothesis testing was adjusted using the Bonferroni method. The corresponding data are presented in Supp Figure 9. Significance levels: ns, not significant; ****, p-value < 0.0001.

| Group1 | Mean1 | SD1 | n1 | Group2 | Mean2 | SD2 | n2 | statistic | p | p.adj | significance |
|---|---|---|---|---|---|---|---|---|---|---|---|
| Q344X+DMSO | 0.2383 | 0.1089 | 32 | WT+DMSO | 0.2153 | 0.1097 | 32 | 570 | 0.44 | 1 | ns |
| Q344X+DMSO | 0.2383 | 0.1089 | 32 | Q344X+AMI | 0.4075 | 0.1510 | 32 | 193 | 1.90E-05 | 9.50E-05 | **** |
| Q344X+DMSO | 0.2383 | 0.1089 | 32 | Q344X+MAP | 0.2341 | 0.1202 | 32 | 512 | 1 | 1 | ns |
| Q344X+DMSO | 0.2383 | 0.1089 | 32 | Q344X+DIF | 0.4707 | 0.1172 | 32 | 70.5 | 3.19E-09 | 1.60E-08 | **** |
| Q344X+DMSO | 0.2383 | 0.1089 | 32 | Q344X+PRE | 0.2779 | 0.1470 | 32 | 410.5 | 0.175 | 0.875 | ns |
